# Direct control of translational elongation by TAOK2β highlights altered protein synthesis as a fundamental underlying component of autism

**DOI:** 10.1101/2022.08.22.504812

**Authors:** Melad Henis, Tabitha Rücker, Robin Scharrenberg, Melanie Richter, Lucas Baltussen, Durga Praveen Meka, Birgit Schwanke, Nagammal Neelagandan, Danie Daaboul, Nadeem Murtaza, Christoph Krisp, Sönke Harder, Hartmut Schlüter, Matthias Kneussel, Irm Hermans-Borgmeyer, Joris de Wit, Karun K. Singh, Kent E. Duncan, Froylan Calderón de Anda

## Abstract

Microdeletions in the *16p11.2* region of the human genome are frequently associated with autism spectrum disorders (ASDs), but how these genomic rearrangements cause ASD remains unclear. Here, we reveal that TAOK2β, a protein isoform encoded by the human *TAOK2* gene located in the *16p11.2* locus, regulates mRNA translation. To identify key functional interaction partners of TAOK2β, we performed proteomic screening from Neuro-2a (N2a) cells, mouse cortices, and cultured neurons. This revealed translation factors as a major class of enriched interacting proteins. Consistently, TAOK2β is present in mouse cortical polyribosomes and cortices from *Taok2* knockout mice show increased ribosome density on mRNAs and enhanced protein synthesis. Several lines of evidence support an effect of TAOK2β on translation elongation via phosphorylation of eukaryotic elongation factor (eEF2). TAOK2 can directly phosphorylate eEF2 on Threonine 56 and this phosphorylation is reduced in cortices from *Taok2* knockout mice. TAOK2β WT overexpression increased eEF2 phosphorylation levels and reduced protein synthesis, whereas a kinase-dead allele of TAOK2β showed opposite effects. Finally, we show that cortices from the mouse model of the human *16p11.2* microdeletion have increased polysome/monosome (P/M) ratios and protein synthesis, phenocopying *Taok2* loss of function. Importantly, defective translation phenotypes observed in the mouse *16p11.2* microdeletion model of ASD could be normalized either by reintroducing *Taok*2 in vivo or by delivering TAOK2β to cortical neurons derived from *16p11.2* microdeletion mice. Our results uncover a critical role of TAOK2β as a regulator of protein synthesis and support the idea that translational control is a common endpoint of ASD-associated signaling pathways.

## Introduction

ASDs are a group of neurodevelopmental disorders (NDDs) with varying symptoms and severity, for which there is no single known cause. ASDs affect a high proportion of the human population, with an incidence of 1 in 54 children in the US ^1^. Common symptoms in ASD individuals are impaired social interaction and cognition, and repetitive behaviors^2^. Genetic studies have shown an association between copy number variations (CNVs) and neurodevelopmental conditions, including ASDs ^3^. Such associations were frequently described for the human *16p11.2* locus, which contains 31 genes ^4^. The *16p11.2* microdeletion is linked to ASDs and contributes to approximately 1% of all diagnosed cases of ASD ^5^. In contrast, reciprocal microduplications of the *16p11.2* led to schizophrenia ^6^. However, it remains unclear, which of the ∼30 genes localized in the *16p11.2* region are causally linked to behavioral, functional, and anatomical changes observed in ASDs.

The human *TAOK2* gene is localized in the *16p11.2* chromosomal region and has been implicated in neurodevelopmental disorders ^5^. TAOK2 is a serine/threonine kinase with two main spliced isoforms, the long TAOK2α isoform with 1235 amino acids and the shorter TAOK2β with 1049 amino acids. TAOK2-α/β isoforms share exons 1-16 (same residues 1 to 745) but differ at their C-termini ^7, 8^. Whole-genome and exome sequencing of ASD families identified novel *de novo* and inherited mutations in both TAOK2 isoforms^9^. A Taok2 deficient mouse model has abnormalities in brain morphology and irregularities in synaptic and dendritic morphology ^9^. Moreover, these knockout mice showed abnormal postnatal hippocampal-prefrontal brain connectivity, impaired cognition and social behavior depending on *Taok2* dosage during adolescence ^9^.

Whether TAOK2 protein isoforms contribute differentially to the processes affected by altered *TAOK2* genetic dose is still not fully understood. Here, we show that TAOK2β interacts with the translational machinery and is present in polyribosome-mRNA translation complexes in cultured neurons and developing mouse brains. Moreover, we found that genetically altering TAOK2 levels affected translation, with protein synthesis increased in the absence of TAOK2β and reduced by TAOK2β overexpression, suggesting a previously undescribed function of TAOK2β as a translational repressor. This role requires the kinase activity of TAOK2β, and we found that TAOK2β interacts with eEF2 and modulates its phosphorylation status. Cortices from the *16p11.2* deletion mouse model of ASD showed increased P/M ratios as well, suggesting altered translation in vivo in this ASD model. Accordingly, cultured cortical neurons derived from the *16p11.2* deletion mouse model showed exaggerated protein synthesis, which was rescued after reintroduction of TAOK2β. Taken together, our results support the unifying theory that translational control is a common endpoint of ASD-associated pathways ^10, 11^, and improve our understanding of how the *16p11.2* deletion contributes to ASD.

## Results

Previously, we reported that, while TAOK2α is associated with microtubules, TAOK2β has a functional association with the actin cytoskeleton through binding to and affecting RhoA activity ^9^. However, the role of TAOK2β in neuronal differentiation is still not well understood. Here, we used a complementary combination of open-ended screening approaches to gain further insight into the functions of TAOK2β in the developing nervous system. First, we performed immunoprecipitation coupled with mass spectrometry (*IP-MS*) analysis using an anti-TAOK2β antibody in cytoplasmic lysates from either mouse cortices or N2a cells transfected with wild-type TAOK2β. In parallel, we used a neuronal proximity-based proteomic system to identify protein-protein interaction (*PPI*) networks associated with TAOK2β in mouse cortical neurons (18DIV) infected with lentiviral constructs, expressing a BioID2 fusion protein (pLV-hSyn-tGFP-P2A-TAOK2β-13xLinker-BioID2-3xFLAG) ^12^. Co-purified proteins from the three biological samples were involved in chloride channel activity, synaptic structure constituents and translational control based on gene ontology (GO) term analysis (Figure 1 and Supplementary Table 1). Moreover, a detailed analysis of potential translational involvement of TAOK2β revealed connections to RNA processing, translation initiation, and translation elongation (Supplementary Figure 1).

**Figure 1:**
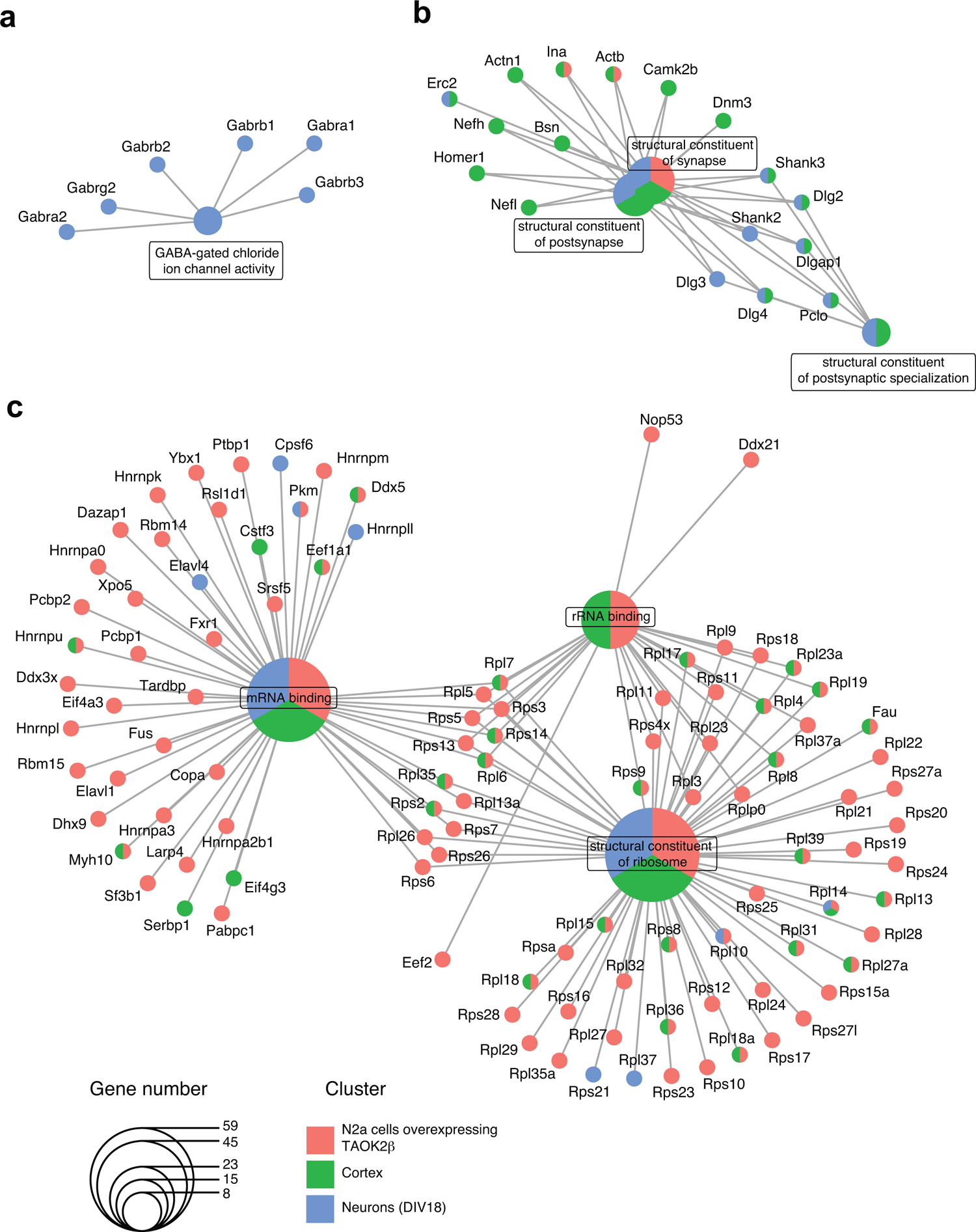
TAOK2β associates with proteins involved in translational regulation, neuronal synaptic structure, and GABA-gated chloride ion channel activity. Overrepresentation analysis as cnetplot of the filtered MS-detected proteins that were co-purified by an anti-TAOK2β antibody from lysates of N2a cells overexpressing TAOK2β (red), mouse cortex (green) and DIV18 cortical neurons (blue) shows GO enrichments for GABA-gated chloride ion channel activity (a) and structural constituents of synapses (b). GO enrichment for mRNA binding, rRNA binding and structural constituents of the ribosome (c), are all functionally related to translational regulation. Circle size indicates the number of genes associated with the respective GO enrichments (details in Supplementary Table 1), and color indicates the respective biological samples used for GO enrichments.

To investigate whether TAOK2β associates with polyribosome translation complexes, we first performed polysome profiling from mouse cortices. Proteins were extracted from collected gradient fractions and used for immunoblotting analysis with antibodies against TAOK2β and control proteins. This showed the presence of TAOK2β in polysomes of wild-type mice and its absence in polysomes of *Taok2* knockout mice (Figure 2a). The detected TAOK2β signal across the translationally active polysome-associated mRNAs (heavy polysomes) was coincided with the distribution of the cytoplasmic Poly(A)-binding protein (PABP) and large ribosomal subunit protein RPL7a, which served as positive controls. To confirm the specific association of TAOK2β with polysomes, the cytoplasmic lysates from wild-type mouse cortices were treated with the Mg^2+^-chelating agent EDTA to disrupt polysomes before loading on the sucrose gradient. The EDTA-treated profile showed the disappearance of polysomes due to the dissociation of ribosomes from the polyribosome complex following treatment. In contrast, the peaks of monosomes (the 80s) and ribosomal subunits 40s and 60s were increased (Supplementary Figure 2a). Immunoblot analysis of the fractionated gradients from EDTA-treated profiles revealed a clear shift of TAOK2β signal from the heavier polysome fractions to the lighter fractions (unbound ribosomal subunits and monosomes). The shift of TAOK2β coincided with that of PABP1 and RPL7a toward the lighter gradient fractions upon EDTA treatment suggests the presence of TAOK2β in polysomal fractions. This reflects ribosome association rather than spurious co-sedimentation with large, non-ribosomal complexes (Supplementary Figure 2b, c). Altogether, these data demonstrate the specific association of TAOK2β with cytoplasmic ribosome-mRNA translational complexes.

**Figure 2:**
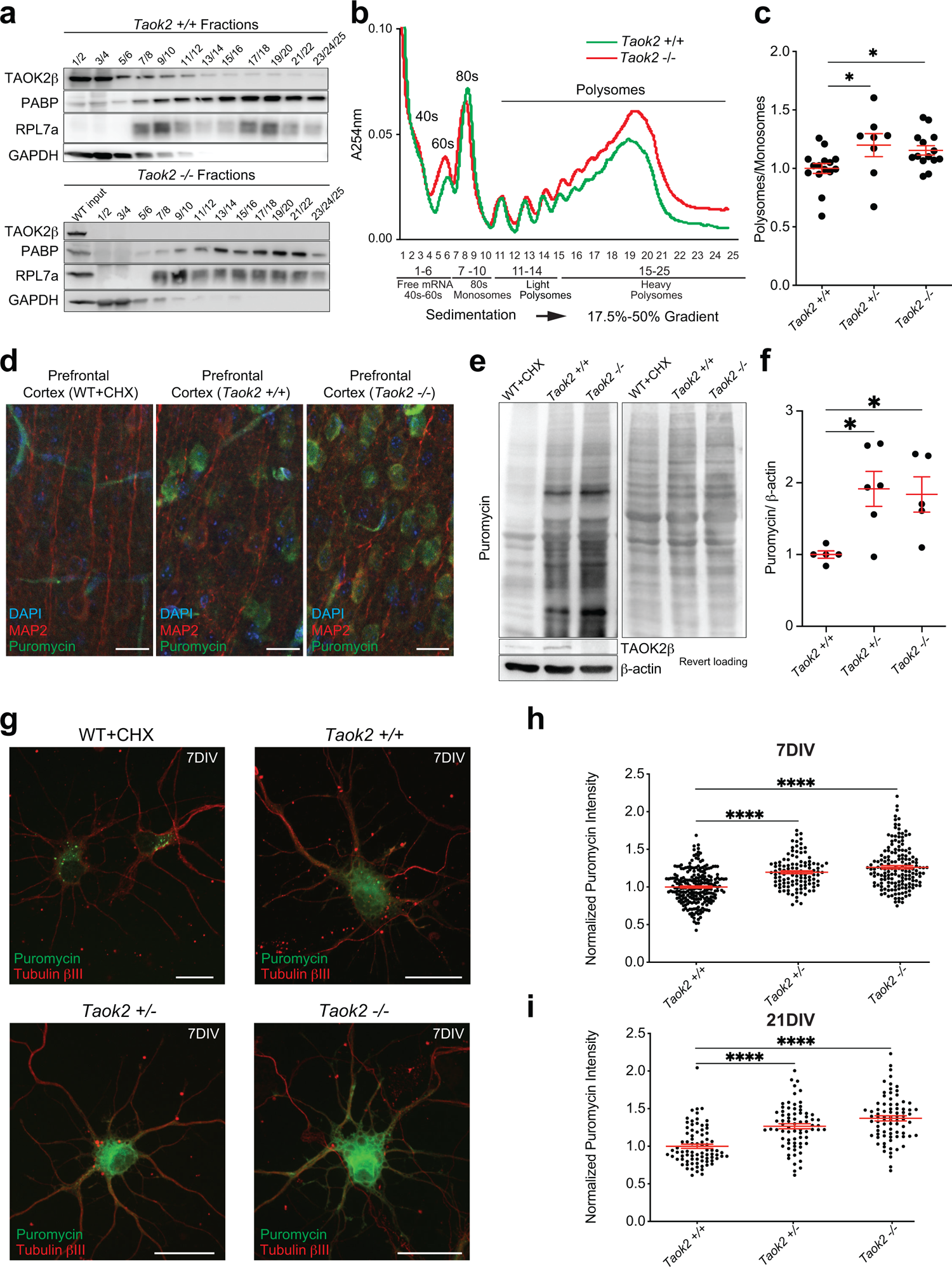
TAOK2β associates with the cytoplasmic polyribosome complex, and its deficiency enhances general translation and protein synthesis. **(a)** Immunoblots of cortical polysome fractions pooled as indicated in the scheme (b), showing the presence or absence of TAOK2β across all polysome fractions isolated from *Taok2* +/+ (upper) or *Taok2* -/- (lower) mouse cortex, respectively. PABP1 and RPL7a were used as positive controls and GAPDH as a negative control to prove the efficiency of polysome preparation. Input from wild-type (WT) cortex verifies the specificity of the TAOK2β antibody. **(b)** Overlay of polysome profiles from *Taok2* +/+ and *Taok2* -/- mouse cortices reveals an increase in the P/M ratio in cortex lacking Taok2. Bottom: Scheme indicates different polysome profile components aligned to their corresponding fraction numbers that sedimented on 17.5-50% sucrose gradients. **(c)** Quantification of polysome profiles shows increased P/M ratios in *Taok2* -/- and *Taok2* +/- mice cortices compared to cortices from *Taok2* +/+ mice. Number of animals, *Taok2* +/+ n = 13, *Taok2* +/- n = 8, *Taok2* -/- n = 12; SEM error bars, *P < 0.05; ordinary one-way ANOVA followed by Tukey’s multiple comparisons test. **(d)** Images of acute cortical slices from the prefrontal cortex of *Taok2* +/+ and *Taok2* -/- mice. Slices were incubated with puromycin to label newly synthesized proteins and immunostained with anti-puromycin (green), anti-MAP2 (red) antibodies, and DAPI (blue) to show increased puromycin fluorescence intensity in Taok2 deficient cortex. Wild-type slices pre-treated with the translation inhibitor cycloheximide (WT+CHX) before puromycin treatment verifies the specificity of the puromycin antibody signal. **(e)** Immunoblot from acute slices analyzed by SUnSET assay shows increased newly synthesized proteins in Taok2 deficient cortex using an anti-puromycin antibody. Revert loading control was used to show equal protein loading for immunoblot (right). **(f)** Quantification for the immunoblot SUnSET assay of cortical acute slices shows the increase of densiometric signal of puromycin normalized to β-actin in *Taok2* -/- and *Taok2* +/- mice. Number of animals, *Taok2* +/+ n = 5, *Taok2* +/- n = 6, *Taok2* -/- n = 5; SEM error bars, *P < 0.05; ordinary one-way ANOVA followed by Tukey’s multiple comparisons test. **(g)** Immunostainings of primary neurons (7DIV), treated with puromycin in the SUnSET assay, immunostained with anti-puromycin (green) and anti-tubulin βƖƖƖ (red) antibodies show increased protein synthesis in Taok2 deficient neurons. Cultured wild-type neurons were pre-treated with cycloheximide (WT+CHX) before puromycin treatment to verify the specificity of the puromycin antibody signal. **(h**, **i)** Quantifications of immunofluorescent SUnSET assay of 7DIV neurons or 21DIV, respectively, show increased normalized puromycin labelling intensity in Taok2 deficient neurons. Number of 7DIV cells, *Taok2* +/+ n= 236, *Taok2* +/- n= 113, *Taok2* -/- n= 166, from 5 different individual embryos for *Taok2* +/+ and 4 embryos for each *Taok2* +/- and *Taok2* -/-. Number of 21DIV cells, *Taok2* +/+ n = 81, *Taok2* +/- n = 73, *Taok2* -/- n = 79, from 3 different individual embryos for *Taok2* +/+ and *Taok2* -/-, and 2 embryos for *Taok2* +/-. SEM error bars, **** p < 0.0001, ordinary one-way ANOVA followed by Tukey’s multiple comparisons test.

To examine whether the presence of TAOK2 in polyribosome complexes is isoform-specific, we transiently transfected N2a cells with plasmids expressing the TAOK2α or TAOK2β isoform, respectively. Next, polysome profiling was performed and the extracted proteins from the collected polysome fractions were analyzed by immunoblot. Immunoblot analysis revealed the presence of both α and β isoforms in the collected polysome fractions across the gradient (Supplementary Figure 2d, e). However, the Myc-tag signal of TAOK2α was less pronounced in heavy polysomes compared with the Myc-tag signal of TAOK2β that localized in light and heavy polysomes. Moreover, we determined the amount of each Taok2 isoform in polysomes from mouse cortices. To this end, we performed liquid chromatography with mass spectrometry coupling (LC-MS/MS) analysis of the extracted proteins from polysomes. Accordingly, we found that TAOK2β was more abundant in polysomes (65.21%) than TAOK2α (34.79%) (Supplementary Figure 2f and Supplementary Table 2) suggesting that a significant fraction of TAOK2β may be involved in regulation of translation.

Next, we aimed to assess the effect of *Taok2*-dependent regulation of translation in brain tissues. Therefore, we performed polysome profiling from cortices of *Taok2* deficient mice (*Taok2* -/- and *Taok2* +/-). Polysome/monosome (P/M) ratios, which indicate the cellular translational state, revealed that the absence of *Taok2* leads to significant changes in global translation. Cortices from *Taok2* deficient mice exhibited increased P/M ratios, strongly suggesting enhanced translation in the *Taok2* deficient mice (Figure 2b, c).

Changes in ribosome density in polysome assays imply effects on the translational machinery, but do not necessarily reflect corresponding directional effects on protein synthesis. For example, increased ribosome density could be due to more ribosomes actively translating, enhanced ribosome stalling, or failure to clear defective ribosomes via quality control mechanisms ^13, 14^. To distinguish between these scenarios and gain more insight into how TAOK2 affects protein synthesis, we used the surface sensing of translation “SUnSET” assay ^15, 16^ to measure the global protein synthesis in acute brain slices from 4-week-old mice. Consistent with an increased P/M ratio from the polysome profiling, puromycin labeling of newly synthesized proteins was also increased in brain slices from *Taok2* -/- and *Taok2* +/- mice as compared to *Taok2* +/+ mice (Figure 2d-f). To determine whether protein synthesis is enhanced in neurons, we cultured cortical neurons from *Taok2* -/-, *Taok2* +/- and *Taok2* +/+ mice. At DIV7 or DIV21, cultured neurons were treated with puromycin to label the newly synthesized proteins, and quantitative immunofluorescent SUnSET analysis was performed. Consistent with what we found in cortical slices, *Taok2* -/- and *Taok2* +/- neurons showed a significant increase in the incorporated puromycin signal at the soma compared to the control *Taok2* +/+ neurons (Figure 2g-i), reflecting increased neuronal protein synthesis. Moreover, gene set enrichment analysis (GSEA) of the *Taok2* -/- total proteome in different brain regions confirmed our previous findings that the cytoplasmic translation is affected, with many ribosomal proteins being significantly upregulated throughout the brain (Supplementary Figure 7b, c and Supplementary Table 3). These data demonstrate that Taok2 is a repressor of translation in developing neocortical neurons.

We next asked whether the physical association of TAOK2β with mRNA-ribosome translation complexes reflects a functional role in regulating translation. To this end, we expressed mutated loss-of-function TAOK2β and TAOK2α (containing an A135P mutation, a human *de novo* missense mutation, which renders this protein into a kinase-dead form of TAOK2 ^9^) in N2a cells and evaluated their polysome profiles. Quantification of polysome profiling assays revealed that the P/M ratios were significantly increased in cells expressing TAOK2β A135P relative to control cells but were not affected by TAOK2α A135P expression (Supplementary Figure 3a-f). This suggests that there are more ribosomes bound to cellular mRNAs after introducing the TAOK2β kinase-dead protein. Conversely, the P/M ratio was decreased after the introduction of wild-type TAOK2β in N2a cells (Supplementary Figure 3g-l). These data point to a functional connection between TAOK2 kinase activity and translation regulation. Moreover, given that the A135P mutation leading to defective kinase activity was originally identified in an ASD patient ^9^, it suggests that defective translation control could increase susceptibility ASD.

Furthermore, SUnSET assay analysis ^16^ of N2a cells either transfected with wild-type TAOK2β or with TAOK2β A135P showed decreased or increased levels of puromycin incorporation into the newly synthesized proteins, respectively (Supplementary Figure 4a-f). Finally, we used ASD patient-derived lymphoblastoid cell line (LCL) expressing *de novo* mutations in TAOK2: TAOK2 A135P (present in both isoforms) and a C-terminal deletion (P1022*), present only in the TAOK2β isoform, which renders TAOK2β into an unstable protein ^9^. Accordingly, polysome profiling and protein synthesis analysis from patient derived LCLs with the A135P mutation revealed increased P/M ratios (Supplementary Figure 5a, b) together with increased levels of newly synthesized proteins, as determined by immunoblot of puromycin (Supplementary Figure 5c, d), compared to the cells derived from the non-affected father. Similarly, the cells of the patient with the C-terminal P1022* deletion have both increased P/M ratios by polysome profiling analysis and increased global protein synthesis by SUnSET assay, as compared with the non-affected father (Supplementary Figure 5e-h). Likewise, N2a cells expressing TAOK2β P1022* mutation showed increased P/M ratios by polysome profiling analysis (Supplementary Figure 6a-c). Altogether, these results demonstrate that TAOK2β regulates translation as a repressor of protein synthesis in N2a cells and ASD patient derived LCL cells.

Global protein synthesis is regulated by many different processes, but especially by phosphorylation of translation factors, ribosomal proteins, and other supporting proteins (e.g. RNA-binding proteins) ^17^. Given that TAOK2β is a kinase and we found that its kinase activity was required for normal regulation of protein synthesis, we hypothesized that altered phosphorylation of specific translation machinery components or regulators would underlie the effects on translation. Thus, to understand how TAOK2β exerts its function as a repressor of translation, we used a labelled phosphoproteomics approach (Supplementary Figure 7a) in multiple brain regions of *Taok2* +/+, *Taok2* +/- and *Taok2* -/- mice to determine the potential phosphorylation targets of Taok2. Identified phosphorylation sites were filtered for the minimal consensus motif of Taok2 (Yadav et al. 2017) to select for potential direct substrates of its kinase activity. High-confidence targets were expected to be significantly downregulated in at least two *Taok2* -/- brain regions. Three phosphorylation sites met these requirements: LPPR3 Thr374, CRMP1 Thr509, and eEF2 Thr57 (Figure 3a and Supplementary Table 4).

**Figure 3:**
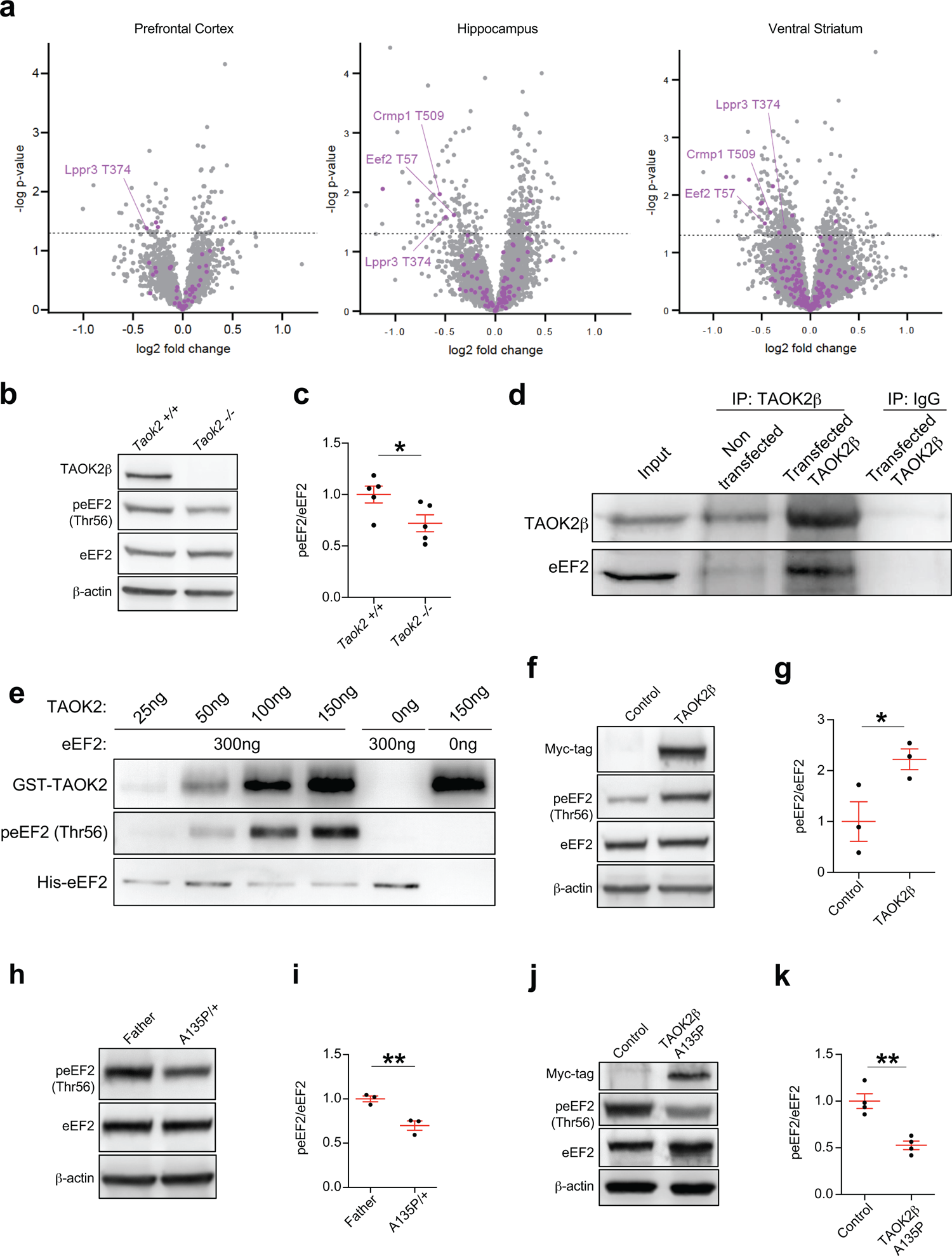
TAOK2β binds eEF2 and represses translation through direct phosphorylation of its Thr56 residue. **(a)** Volcano plot of *Taok2* knockout / *Taok2* wild-type phosphoproteome show potential direct substrates of Taok2 concurringly identified in three different brain regions (left: Prefrontal Cortex, middle: Hippocampus, right: Ventral Striatum) and downregulated in *Taok2* knockout brains. Phosphopeptides containing the TAOK2 motif pTXX[K/R/H] are in purple. Two data points are outside of axis limits. **(b, c)** Immunoblot and quantifications of cortical lysates from *Taok2* +/+ and *Taok2* -/- mice show significantly decreased phosphorylated eEF2 (Thr56) normalized to total eEF2 levels in cortices of *Taok2* -/- animals. Number of animals per each genotype n=5; *P < 0.05, SEM. error bars, unpaired t-test. β-actin was used as a loading control. **(d)** Immunoblot of the TAOK2β immunoprecipitation (IP) from non-transfected N2a cells and N2a cells overexpressing wild-type TAOK2β shows the interaction of TAOK2β with eEF2. Unspecific rabbit IgGs serve as IP controls and total cytoplasmic N2a cell lysate (input) verifies the specificity of the antibodies. **(e)** Immunoblot of ascending amounts of a recombinant GST-tagged TAOK2 (aa 1-314) protein incubated with a recombinant His-tagged eEF2 or controls (150 ng TAOK2 without eEF2 or 300 ng eEF2 protein without TAOK2) used in an *in vitro* kinase assay as indicated. Direct phosphorylation of Thr56 of eEF2 protein by the increased concentration of TAOK2 protein was detected by immunoblotting analysis using an anti-phospho-Thr56 specific antibody. **(f, g)** Immunoblot and quantifications of N2a cell lysates transfected with wild-type TAOK2β and its respective control show significantly increased ratios of the phospho-eEF2 compared to total eEF2 in cells transfected with wild-type TAOK2β. Number of biological replicates: n=3 per condition from different independent transfection experiments; *P < 0.05, SEM. error bars, unpaired t-test. β-actin was used as a loading control. **(h, i)** Immunoblot and quantifications of LCLs lysates with TAOK2 mutation A135P at its kinase domain and non-affected father show significantly decreased ratios of the phospho-eEF2 compared to total eEF2. Number of biological replicates: n=3 per condition; **P < 0.05, SEM. error bars, unpaired t-test. β-actin was used as a loading control. **(j, k)** Immunoblot and quantifications of N2a cell lysates transfected with TAOK2βA135P and its respective control show significantly decreased ratios of phospho-eEF2 compared to total eEF2 in cells transfected with TAOK2βA135P. Number of biological replicates: n=4 per condition from different independent transfection experiments; **P < 0.01, SEM. error bars, unpaired t-test. β-actin was used as a loading control.

The phosphorylation status of elongation factor eEF2 is crucial in regulating translation at the elongation step, impacting on ribosome translocation along the mRNA^18^. Especially phosphorylation of residue Thr56 (corresponding to mouse Thr57) is known to strongly inhibit eEF2 activity and thereby the rate of translation ^19^. Therefore, we validated the finding that eEF2 Thr57 phosphorylation is reduced in *Taok2* -/- mouse brains by using a well-established rabbit anti-phospho-eEF2 (Thr56) antibody in mouse cortical lysates (Figure 3b, c). This phosphorylation is usually ascribed to the eukaryotic elongation factor 2 kinase (eEF2K) and has already been linked to Fragile X syndrome (FXS) ^20^. Similarly, phosphorylation of eEF2 by TAOK2β could act as a potent elongation inhibitor in global protein synthesis.

According to our initial IP-MS approach, we also detected eEF2 in protein complexes immunoprecipitated with the TAOK2β antibody (Figure 3d), confirming the result obtained after overexpression of TAOK2β in N2a cells (Figure 1 and Supplementary Table 1). To determine whether TAOK2 can directly phosphorylate eEF2, we incubated bacterially purified His-tagged eEF2 protein with human purified GST-tagged TAOK2 kinase domain (aa 1-314) in an *in vitro* kinase assay. We found that in the absence of other proteins, small amounts of TAOK2 can already lead to phosphorylated eEF2 Thr56, indicating this is directly mediated by TAOK2 (Figure 3e). Similarly, overexpression of TAOK2β in N2a cells causes increased phosphorylation of eEF2 (Figure 3f, g) concomitant with decreased amounts of newly synthesized protein, assessed by puromycin labeling (Supplementary Figure 4b, d-f). On the contrary, we observed reduced phosphorylation of eEF2 (Thr56) and increased levels of newly synthesized protein in ASD patient derived LCLs containing the TAOK2 A135P mutation (Figure 3h, i and Supplementary Figure 5c, d) and in N2a cells expressing TAOK2β A135P (Figure 3j, k, and Supplementary Figure 4c, d). Moreover, we could not detect phosphorylation changes in the only known upstream kinase of eEF2, eEF2K ^19^ (Supplementary Table 4 and Supplementary Figure 8), further suggesting that TAOK2 directly phosphorylates eEF2 rather than functioning upstream of eEF2K. Altogether, these results suggest that TAOK2β kinase activity inhibits eEF2 activity via phosphorylation of its Thr56 residue leading to decreased protein synthesis.

Given that in humans *TAOK2* localizes in the genomic *16p11.2* NDD susceptibility locus, we next asked whether the mouse model of *16p11.2* deletion (B6;129S7-Del(7Slx1b-Sept1)4Aam/J+/-, abbreviated as *16p11.2del* +/-) recapitulates our findings regarding dysregulated global translation. Thus, we analyzed the polysome profiles of cortices from a *16p11.2del* +/- mice ^21^. We found increased P/M ratios (Figure 4a, b), similar to what we detected for cortices of the *Taok2* -/- mice (Figure 2b, c), reflecting enhanced ribosome density on cellular mRNAs. Accordingly, cultured cortical neurons from the *16p11.2del* +/- mice treated with puromycin showed an elevated rate of puromycin labeling of the newly synthesized protein compared to WT cells (Figure 4c-e). Our results provide clear evidence that TAOK2 is a translation repressor; therefore, we aimed to rescue the effect of enhanced translation in the*16p11.2del* +/- mouse model. We generated a transgenic mouse B6; 129S7-*(Del(7Slx1bSept1)Tg(TaoK2Hhtg*)Cal (abbrev. *delTgTaoK2*), where *Taok2* was re-introduced in the *16p11.2del* +/- mouse model and cortices were analyzed by polysome profiling assay. Interestingly, the increased P/M ratios from *16p11.2del* +/- mouse cortices were normalized to wild type levels by re-introduction of *Taok2* on a transgene (Figure 4f-h) implying that *TAOK2* alone is likely to mediate the impact of *16p11.2* deletion on translation. Finally, we tested whether the TAOK2β alone could rescue the exaggerated protein synthesis phenotype in *16p11.2del* +/- neurons. To this end, we used *in utero* electroporation (IUE) to transfect *16p11.2del* +/- cortices with TAOK2β and tDimer at embryonic day 14.5-15 (E14.5-E15) and prepared primary cortical neuronal cultures two days later (E17). After seven days *in vitro* (7DIV), cultured neurons were treated with puromycin to label the newly synthesized proteins and prepared for immunostaining using puromycin and TAOK2β antibodies. Notably, after the re-introduction of TAOK2β, the levels of newly synthesized proteins were reduced (Figure 4i, j), confirming the role of TAOK2β as a repressor of protein synthesis.

**Figure 4:**
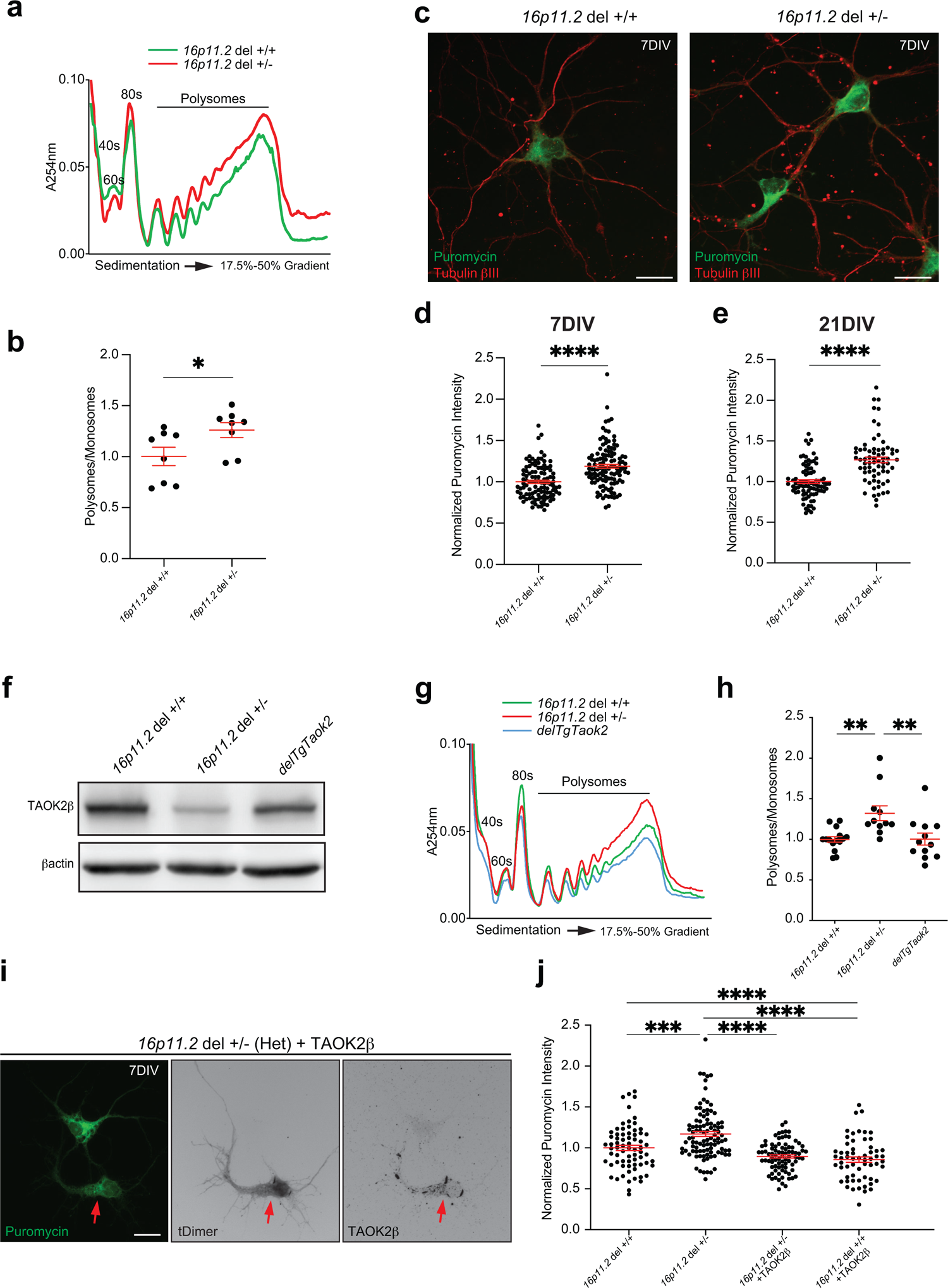
*16p11.2* microdeletion displays dysregulated global translation that is rescued in the presence of TAOK2. **(a)** Overlay of polysome profiles from *16p11.2del* +/+ and *16p11.2del* +/- mice cortices shows an increase in P/M ratio in cortices of *16p11.2del* +/- mice **(b)** Quantification from (a) showing significantly increased P/M ratios in *16p11.2del* +/- mouse cortex in comparison with the wild-type cortex. Number of animals, *16p11.2del* +/+ n =8, *16p11.2del* +/- n = 8; SEM error bars, *P < 0.05; SEM. error bars, unpaired t test. **(c)** Immunostainings of primary neurons 7DIV treated with puromycin and immunostained with anti-puromycin (green) and anti-tubulin βƖƖƖ (red) antibodies show increased puromycin labelling intensity in *16p11.2del* +/- deficient neurons. **(d, e)** Quantifications of normalized puromycin labeling intensity at 7DIV neurons or 21DIV, respectively, show increased newly synthesized protein by SUnSET assay in *16p11.2del* +/- deficient neurons. Number of 7DIV cells, *16p11.2del* +/+ n= 107, *16p11.2del* +/- n= 119, from 3 different individual embryos of *16p11.2del* +/+ and 4 embryos of *16p11.2del* +/-. Number of 21DIV cells, *16p11.2del* +/+ n= 85, *16p11.2del* +/- n= 68, from 5 different individual embryos of *16p11.2del* +/+ and 4 embryos of *16p11.2del* +/-. SEM error bars, **** p < 0.0001, ordinary one-way ANOVA followed by Tukey’s multiple comparisons test. **(f)** Immunoblot shows the expression of TAOK2β in the cortices of *16p11.2del* +/+, *16p11.2del* +/- and *delTgTaoK2* mice. **(g)** Overlay of polysome profiles from *16p11.2del* +/+, *16p11.2del* +/- and *delTgTaoK2* mice cortices shows an increase in P/M ratio in cortices of *16p11.2del* +/- mice and its restoration to the normal wild-type levels in *delTgTaoK2* mice. **(h)** Quantifications of polysome profiles show significantly increased P/M ratios in cortices of *16p11.2del* +/- mice. Number of animals, *16p11.2del* +/+ n =15, *16p11.2del* +/- n = 11, *delTgTaoK2* n=13 (one value was identified as an outlier); SEM error bars, **P < 0.01; SEM. error bars, ordinary one-way ANOVA followed by Tukey’s multiple comparisons test. **(i)** Immunostaining of *16p11.2del* +/- cortical neurons after IUE at E14.5-E15 with TAOK2β and t-dimer2 plasmids treated at 7DIV with puromycin and immunostained with anti-puromycin (green) and anti-TAOK2β (grey) antibodies shows a reduction of puromycin labeling intensity in the transfected cells. **(j)** Quantifications from (i) show a significant increase of the normalized puromycin labeling intensity in *16p11.2del* +/- neurons compared to their corresponding controls. This increased intensity is significantly reduced to the normal wild type levels after transfection with TAOK2β. Number of cells, non-transfected *16p11.2del* +/+ n = 71, *16p11.2del* +/+ + TAOK2β n = 60 from 4 different embryos; non transfected *16p11.2del* +/- n = 99, *16p11.2del* +/- + TAOK2β n = 84 from 4 different embryos. SEM error bars, ***P < 0.001, **** p < 0.0001, ordinary one-way ANOVA followed by Tukey’s multiple comparisons test.

Altogether, our results demonstrate a new role for TAOK2β in translational control of protein synthesis via interaction with the translational machinery and phosphorylation of the elongation factor eEF2. Moreover, we observe the same translation phenotypes in a mouse model for ASD caused by the *16p11.2* microdeletion. TAOK2 appears to be the key mediator of these translation phenotypes, given that reintroducing *TAOK2* transgenically or TAOK2β alone in neurons is sufficient for rescue. Thus, our data implicate altered translational regulation due to the lack of *TAOK2* as an underlying cause of ASDs linked to the *16p11.2* microdeletion.

## Discussion

### The ASD-susceptibility gene *TAOK2* associates with the translation machinery and represses translation

Translational control is important for gene expression in response to different physiological and pathological conditions ^22^. Many ASD-associated genes encode proteins that are components of the translational machinery and regulatory proteins involved in different stages of translation (e.g., eIF4E, eIF3g, eEF1A2, RPL10, eIF4EBP2, and UPF3B) ^23^. Moreover, many RNA-binding proteins (RBPs) which regulate mRNA translation are known ASD risk genes as well. For example, *FMRP*, a well-known ASD-associated gene, is one of the best-characterized RBPs functioning as a translational repressor, and its deficiency underlies the pathogenesis of FXS ^24^.

The strong connections between dysregulated translation factors and ASD have led to the proposal over a decade ago that failure to properly tune the levels of neuronal protein synthesis might be a fundamental cause of ASD ^11, 25, 26^. This idea is supported by more recent work demonstrating a key role for translation initiation regulatory pathways in mediating pathobiology in ASD models, as well as the potential to correct these pathways for therapeutic benefit ^10, 27, 28^. However, it has remained unclear if 16p11.2 CNV-mediated ASD connects to translation; no gene in the locus encodes a protein previously implicated in protein synthesis or its regulation. Here, we show for the first time that a gene located in the *16p11.2* ASD susceptibility locus is associated with translational control. Specifically, we reveal that TAOK2 can act as a repressor regulating protein synthesis at the elongation step. Previously, different proteins involved in translation were among numerous proteins identified as potentially enriched binding partners of Taok2 using a peptide pull-down method from neuronal lysate ^29^. Additionally, our protein screen and analysis of purified polyribosome complex proteins using Taok2-specific antibodies confirmed the presence of Taok2 protein in polyribosome translation complexes. Presumably interaction with ribosome-mRNA translation complexes is mediated through one or more protein-protein interactions rather than interaction with the mRNA, since Taok2 and its human ortholog both lack a known RNA binding motif, and it was not identified as a non-canonical mRNA-binding protein in the global mRNA interactome capture experiments ^30, 31^. Translation involves dynamic interactions between multiple-subunit initiation factors, RNA-binding proteins, and large protein-RNA complexes (i.e., the ribosome). We capture most of these in our pulldowns and in principle any of them might recruit TAOK2. Regardless of how TAOK2 is recruited to the translational machinery, these data suggested a functional role for TAOK2 in regulation of translation, which we could substantiate via multiple lines of evidence.

A role for any 16p11.2 encoded gene in translation is particularly interesting, since several studies suggest that disturbances in the translational control machinery and the subsequent dysregulation of synaptic proteins are one of the molecular mechanisms underlying the etiology for synaptopathies in ASD ^10, 11^. Therefore, dysregulated protein synthesis is considered a common feature in most autistic phenotypes, and excessive neuronal protein synthesis is observed in particular in FXS pathogenesis ^32^ and in other forms of autism ^10, 27^. In this regard, measurement of the amount of newly synthesized protein by the SUnSET assay showed enhanced protein synthesis in cortex and neurons from *Taok2* deficient mice compared to the wild-type controls. These data suggest that Taok2 is a repressor of neuronal translation, like many other known autism-linked genes, such as *FMRP*, *PTEN*, and *TSC1/2*. Mutations in these genes disturb translation repression, cause aberrant protein synthesis and lead to the autistic phenotypes ^33–35^. Consistently, analysis of *de novo* global protein synthesis in animal models of FXS and in different cell types derived from FXS patients, such as lymphoblastoid cells, fibroblasts as well as an induced pluripotent stem cell (iPSC)-derived neural progenitors (NPCs), revealed the increased global protein synthesis due to the absence of translation repression by *FMRP* ^36–38^.

Similarly, we found that lymphoblastoid cells from autistic patients with TAOK2 mutated at the residue A135P in the kinase domain, which leads to loss of TAOK2 kinase function, showed enhanced ribosome density on mRNAs by polysome profiling analysis and a corresponding increased amount of newly synthesized proteins compared to cells derived from the non-affected father. These data indicate that the role of TAOK2 in translational repression is mainly achieved through its kinase function. Furthermore, N2a cells, a neural crest-derived cell line, transfected with wild-type TAOK2β revealed reduced translation compared to the control. However, when these cells were transfected with the kinase dead TAOK2β A135P mutation, global translation analyzed by polysome profiling and the amount of protein synthesis measured by puromycin incorporation, were elevated in comparison with control cells. In fact, cells expressing the TAOK2α A135P mutated isoform did not reveal any significant change in the global translation, as the P/M ratios were similar to the control. Thus, our data suggest that the defects in translational control and global protein synthesis directly result from the loss of TAOK2β kinase function. Its relative abundance in polyribosomes is one possible explanation for the isoform-specific role of TAOK2β because there is no difference in the kinase domain of both proteins. A greater proportion of TAOK2β than TAOK2α appeared to be stably associated with polyribosomes according to LC-MS/MS analysis of extracted proteins from polysomes of wild type mouse cortices and by immunoblot analysis of polysomes from N2a cells transfected with wild type TAOK2β or wild type TAOK2α. Furthermore, the difference in the C-termini of both isoforms ^8, 9^ might affect their association with polyribosomes differentially. Additionally, the auto-regulatory function of the C-terminus ^39, 40^ can affect the translational regulation as observed for the P1002* mutation at the C-terminus (specific for the TAOK2β isoform) in patient-derived LCLs with P1002* mutation and N2a cells expressing TAOK2βP1002* (Supplementary Figure 5e-h and Supplementary Figure 6). Taken together, our data demonstrate that TAOK2 physically associates with the translational machinery to modulate translation as a repressor. Our data further highlight the kinase activity as crucial for TAOK2 function in translational control.

### Taok2 represses translation elongation via direct phosphorylation of eEF2

Protein kinases other than TAOK2 have long been known to phosphorylate different components of the translational machinery, especially the initiation and elongation factors^17^. These phosphorylation events regulate the activity and interactions of these factors with other translational machinery components to thereby regulate translation ^17^. Previous work has implicated known phosphoregulatory pathways that modulate activity of translation initiation factors in in different neuropsychiatric disorders ^27, 41^. Our work suggests that elongation can also be targeted and identifies TAOK2 as a new direct regulator of eEF2. Previous work identified a single eEF2 kinase (eEF2K) that can phosphorylate eEF2 on its Thr56 residue to inhibit its role in ribosome translocation during translation elongation^19, 42, 43^. eEF2K is a Ca^2+^/CaM-dependent kinase with multiple regulatory phosphorylation sites that affect its activity in response to different signalling pathways ^44^. eEF2K activity is inhibited due to phosphorylation of the inhibitory sites: Ser78, Ser366 ^45^, and Ser359 ^46^ mediated via mTORC1 signalling; and Ser359 and Ser366 mediated via the MAP kinase pathway ^45, 47^. Conversely, phosphorylation of Ser500 by cAMP ^48^ and Ser398 by AMPK ^49^ stimulates eEF2K activity. TAOK2 is a serine/threonine kinase, and we did not detect any phosphorylation changes in eEF2K regulatory sites by phosphoproteomics (Supplementary Table 4) or western blot using a phospho-Ser366 specific antibody (Supplementary Figure 8) in our search for TAOK2 targets. This suggests that Taok2 does not directly or indirectly affect eEF2K activity. Furthermore, we showed that the Thr56 residue of eEF2 is a phospho-site modulated by Taok2 and confirmed direct phosphorylation at this site by TAOK2 kinase using an *in vitro* kinase assay. Thus, our study identifies TAOK2 as a novel kinase other than eEF2K that can directly modulate phosphorylation of Thr56 of eEF2. We thereby suggest a new mechanism for regulation of translation elongation and tuning of neuronal protein synthesis via the activity of TAOK2β.

Our data suggest that TAOK2 controls translation via the elongation factor eEF2, however we acknowledge that other mechanisms might also contribute. For example, differential expression protein analysis for different brain regions from *Taok2* knockout mice showed significantly upregulated proteins of the translational machinery components, especially ribosomal proteins (Supplementary Figure 7c and Supplementary Table 3). Upregulation of translation factors, including ribosomal proteins, has been reported in ASD-patient derived neuronal precursor cells and post-mortem cortical tissues and iPSC-derived neural progenitor cells, as well as in the non-neuronal cell types such as lymphocytes from ASD patients ^50–52^. Thus, upregulation of ribosomal proteins is a feature of ASD and might also contribute to in some way to the translation phenotypes that we observe. Although we did not find evidence for altered phosphorylation of known regulators of initiation (4EBPs or eIF2α) in our phosphoproteomics studies, an additional effect of TAOK2 on translation initiation is nevertheless possible in principle. However, since we find effects on eEF2 phosphorylation at specific sites previously determined to impact translation elongation, the simplest interpretation of our data is that this mechanism contributes significantly to the translation phenotypes that we observe in the absence of TAOK2 and its kinase activity.

### TAOK2 is a key mediator of protein homeostasis in *16p11.2* microdeletion ASD models

*TAOK2* is one of ∼30 genes located in the *16p11.2* chromosomal region and has been implicated in the etiology of ASD ^5^. Specifically, the *Taok2* deficient mouse model has similar anatomical and behavioral abnormalities to those identified in *16p11.2* microdeletion mouse models and patients ^9, 53, 54^. In our study, we observed enhanced translation and protein synthesis in the *16p11.2* microdeletion, phenocopying what we observed in a *Taok2* knockout mouse model. Importantly, we were able to normalize the translation phenotypes by the re-introduction of *Taok2* in the *16p11.*2 microdeletion mouse model itself or by direct delivery of TAOK2β to primary neurons derived from that model. Altogether, our data suggest that *TAOK2* is a potential key mediator of protein synthesis abnormalities in carriers of the *16p11.2* CNV. Furthermore, our results highlight previously unappreciated exaggerated protein synthesis due to a lack of TAOK2-mediated translational control in this important genetic model of ASD. Thus, altered translation emerges as a common unifying aspect of ASD, even for genetic variants where the genes themselves do not obviously connect to classical translational control pathways. This suggests that modulation of translation via TAOK2 or other routes might potentially have therapeutic benefit in 16p11.2-caused ASD. Our results also motivate examining whether TAOK2 kinase activity and its impact on eEF2 phosphorylation may be more generally involved in effects on translation in ASD patients outside of the *16p11.2* variant population. Interestingly TAOK2 mRNA is a target, and its regulation is under control of fragile X mental retardation protein (FMRP), making it a high-risk gene associated with CNVs ^55^. Furthermore, FMRP represses translation at the initiation step, where it binds with CYFIP1 and inhibits cap dependent translation initiation by blocking the eIF4E– eIF4G interaction ^56^, as well as at the elongation step by stalling ribosomes on its target mRNAs ^55^. Thus, it will be particularly interesting in the future to investigate whether there are connections between elongation regulation by TAOK2 and FMRP, as well as potential interplay with effects on translation initiation. One could envisage altered translation caused via multiple unlinked routes involving effects on initiation, RBPs, or elongation. However, it seems equally likely that homeostatic mechanisms will involve crosstalk between these different translational regulatory pathways. It will be important to explore these possibilities in future work aimed at understanding the contribution of altered translation to ASD pathobiology and the therapeutic potential for targeting translation to help ASD patients.

## Acknowledgements

We thank Francis Impens and Rupert Mayer (VIB Proteomics Core) for their experimental help. We are thankful to Martina Reiss and Harald Kranz from Gene Bridges GmbH, Heidelberg, Germany (now Gen-H, Heidelberg, Germany) for excellent support and generating the transgenic mouse model B6;129S7-Tg(Del(7Slx1bSept1)TaoK2Hhtg/Cal. L.B. is supported by an FWO Postdoctoral Fellowship (12ZK221N). M.H. is partially funded by a scholarship (ID: seventh plan 2012-2017) from the Cultural Affairs and Missions Sector, Ministry of Higher Education of the Arab Republic of Egypt. NN was partially supported by a Deutsche Akademischer Austauschdienst (DAAD) Predoctoral Fellowship. M.K. is supported by KN556/11-1 (FOR 2419). KED was supported by Deutsche Forschungsgemeinschaft (DFG) Grants: DU 1396/3-1 and DU 1396/4-1; and grant 11/90 from the Werner Otto Stiftung. FCA is supported by Deutsche Forschungsgemeinschaft (DFG) Grants: CA 1495/4-1, and CA 1495/7-1; ERA-NET Neuron Grants (Bundesministerium für Bildung und Forschung, BMBF, 01EW1910, and 01 EW2108B), JPND Grant (Bundesministerium für Bildung und Forschung, BMBF, 01ED1806),

## Material and Methods

### Animals and animal welfare and approvals

*Taok2* -/- mice (B6, *TaoK2^tm1Heb^*/Cal) ^9, 57^ and *16p11.2 del* +/- mice (B6;129S7-*(Del(7Slx1b-Sept1)4 ^AaM^*/Cal, Jax# 013128) ^21^ were bred as previously described. All animals were raised, genotyped, and housed in compliance with standard regulations at the Central Animal Facility at University Medical Center Hamburg-Eppendorf, Hamburg (UKE). All mice experiments were performed according to the German and European Animal Welfare Act. The used experimental procedures have the approval of the Animal Research Ethics Board (AREB), the local authorities of the State of Hamburg (TVA N007/2018), and the animal care committee of UKE.

### Generation of the B6;129S7-(DelSlx1bSept1)4Aam*Tg*TaoK2Hhtg/Cal mouse line

To introduce the mouse *Taok2* gene in the *16p11.2 del* background, heterozygote zygote donor mice were obtained by breeding B6;129S7-(DelSlx1bSept1)4Aam (Jax #: 013128) +/- to wildtype littermates. Superovulation and pronuclear injection were performed as described before ^58^.

The donor plasmid RP23-321H3_bb_Taok2 PB3’Amp_PB5’Kan was generated by Gene Bridges (now Gen-H, Heidelberg, Germany) (Supplemental Figure 9) to allow target integration by PiggyBac transposase. PiggyBac integration sites flank 21.2 kb murine genomic DNA (Chromosome 7: 126,464,301-126,485,468 of GRCm39 reference genome from ENSEMBL) covering the murine Taok2 gene. For further use for a knock-in approach, flanking regions of the mouse locus on chromosome 7 corresponding to the human 16p11.2 locus were recombined at the 5’ and 3’ of the ITR sites. The *Super PiggyBac Transposase* plasmid (System Biosciences; Cat# PB210PA-1) was obtained from BioCat, Heidelberg. Integration of the insert by recombination was confirmed by PCR using 5’-ITR-specific primer PB5-F (GTGCTTGTCAATGCGGTAAGTGTCACTG) with insert-specific primer Taok-rev (GGCATACATTCTTGGAGTCAACTTTACATTGC) and with 3’-ITR-specific primer PB3-rev (GCATGTGTTTTATCGGTCTGTATATCGAGG) with insert-specific primer Taok-F (CACTTTTTGTAATAGGTCTTCTAAACTCAGAAGTG).

To establish the B6;129S7-(DelSlx1bSept1)4Aam*Tg*TaoK2Hhtg/Cal mouse line founder #31 was crossed to B6;129S7+/+ obtained from matings of *16p11.2del* +/- to wild type littermates and offspring was analyzed as described above. Animals carrying the TaoK2 insertion on the *16p11.2del* +/- background were crossed to *16p11.2del* +/- to obtain mice positive for the transgene on *16p11.2del* -/- background.

To confirm the *Taok2* insertion on the *16p11.2del +/-* background animals, genotypes were determined by PCR from mouse-tail DNA using standard protocols. First, the *16p11.2del* genotype was determined using the following primers: forward primer (CCTGAGCCTCTTGAGTGTCC) and reverse primer (GTCGGTTCAGGTGGTAGACG) for the WT allele, forward primer (ACCTGGTATTGAATGCTTGC) and reverse primer (TGGTATCTCCATAAGACAGAATGC) for the mutant allele. Second, the *delTgTaoK2* genotype was determined with the following primers: for the 5’ end, forward primer # 956 PBTP5-FL (GTCCTTGTCAATGCGGTAAGTGTCACTG); reverse primer # 957 TAOK-5’-PBTP-rev (GGCATACATTCTTGGAGTCAACTTTACATTGC); for the 3’ end, forward primer # 1090 TaoK-3-F2 (CACTTTTTGTAATAGGTCTTCTAAACT), and reverse primer # 958 PBTP3-revL (GCATGTGTTTTATCGGTCTGTATATCGAGG).

### Plasmids

Wild-type TAOK2α and β isoforms; and their site directed mutagenesis plasmids that have been described before ^9^ were used for N2a cells transfection in addition to pCDNA3.1-myc (as empty vector control plasmid) and pEGFP-F (plasmid encoding the enhanced GFP sequence with a farnesylated signal). pAAV-CAG huTAOK2β wt (plasmid encodes humanTAOK2-short variant) and pAAV-CAG-tDimer2 (plasmid encodes the A tandem dimer mutant of DsRed tagged to the RFP) were used for *in utero* electroporation of cortical neurons. pLV-hSyn-tGFP-P2A-POI-13xLinker-BioID2-3xFLAG construct expressing BioID2 fusion proteins and Luciferase-P2A-BioID2-3xFLAG construct as a negative control were used in neuron viral infection ^12^.

### Antibodies

The following antibodies were used in these studies for immunocytochemical or western blotting analysis: Rabbit anti-Taok2β, (Synaptic Systems; 395003, WB 1:1,000, ICC 1:250, IP 1:100); mouse anti-puromycin, clone 12D10, (Millipore; MABE343, WB 1:10,000, IHC, ICC 1: 5,000); chicken anti-MAP 2, Synaptic Systems;188006, IHC 1:500); guinea pig anti-beta3-tubulin (Tuj1), (Synaptic Systems; 302304, ICC 1:500); mouse anti-β-Actin, Sigma; A5316, WB 1:2,000); Rabbit anti-β-Actin (13E5), (Cell Signaling Technology; 4970, WB 1:5,000); rabbit anti-Myc-Tag (71D10), (Cell Signaling Technology; 2278, WB 1:1,000, ICC 1:300); rabbit anti-IgG (Millipore, 03-241 IP 1:200); rabbit anti-phospho-eEF2 (Thr56), (Cell Signaling Technology; 2331, WB 1:1,000); rabbit anti-GST-Tag, (Cell Signaling Technology; 2625, WB 1:1,000); mouse anti-His-Tag (27E8), (Cell Signaling Technology; 2366, WB 1:1,000) and mouse anti-eEF2, clone 4B3-G7-H5, (Abcam; ab131202, WB 1:2,000); rabbit anti-phospho-eEF2K (Ser366), (Cell Signaling Technology; 3691, WB 1:1,000); and mouse anti-eEF2K antibody (C-12), (sc-390710; WB 1:200). HRP-(Dianova; WB 1:2,000-5,000) and Alexa Fluor 488 or 647 or 568 conjugated secondary antibodies (Invitrogen; ICC 1:500, IHC 1:300). Nuclei were visualized with Hoechst (4′,6-diamidino-2-phenylindole) (DAPI) (Invitrogen; 3258, ICC/IHC 1:5,000).

### *In utero* electroporation (IUE)

Lateral ventricles of each embryo (E14.5-E15) of pregnant *16p11.2del* +/- mice were microinjected with a combination of TAOK2β (2.5 µg/µl) and tDimer2 plasmids (0.3 µg/µl) as previously described ^9^. Post-surgery, the animals received meloxicam orally (10-20 mg/kg body weight, twice daily) in soft food to relieve post-operative pain until they were euthanized for embryo collection.

### Cell lines

N2a cells were grown in Dulbecco’s modified Eagle’s medium high-glucose GlutaMAX culture medium (Invitrogen), supplemented with 10% fetal bovine serum (Gibco), 1% penicillin/streptomycin antibiotics (Invitrogen) and L-Glutamine 2mM (Invitrogen). N2a cells were transiently transfected with jetOPTIMUS DNA Transfection Reagent (Polyplus, Cat No. 117-15) according to the manufacturer’s guidelines. The total amount of plasmid DNA used for transfection of a 10 cm plate was 9-10 µg (7-8 µg from different TAOK2 variants or empty vector control (pcDNA 3.1) + 2 µg pEGFP-F plasmid). Following 4 hours of transfection, the medium containing the transfection complex was replaced by a pre-warmed fresh medium. 2 hours later, the cells were passaged one time into 10 cm dishes or PPL pre-coated glass coverslips, depending on the experimental conditions. The transfection efficiency was determined directly before harvesting by taking images for GFP expressed signal in control and transfected cells or by immunohistochemical staining using antibodies against the Myc-tag protein or TAOK2β. Only cells with 70-80% transfection efficiency were harvested after 40–48 hours post-transfection and used for experiments.

Lymphoblastoid cell line cells (LCLs) were generated from peripheral blood, mainly mononuclear cells confined from the blood of probands and their families, which were immortalized utilizing the Epstein-Barr virus ^59^. LCLs were grown in a standard suspension condition in RPMI 1640 media (PAN Biotech) supplemented with 15% FBS. 200.000 cells/ml were used for starting a culture in 5 ml, and every 2-3 days, the medium was increased to 20 ml, and the cells were split to avoid over confluency as they are growing in clumps. During the experiments, the cells were sedimented by centrifugation at 1000 r.p.m. for 5 minutes at RT.

### Primary cortical neuron culture

Primary cortical neuron cultures from embryos of pregnant *Taok2* +/- or *16p11.2del* +/- mice were prepared as described previously ^9^. The cells were kept in culture for seven days in vitro (7DIV) or 21 days in vitro (21DIV) at 37°C with 5% CO2 before treatment with puromycin to measure protein synthesis by SUnSET assay. 0.5 μM Cytosine β-D-arabinofuranoside (Ara-C) was added to the cultured cells on the 7^th^ day in vitro (7DIV) to the cells maintained up to 21DIV to inhibit the growth of the glia cells. Cortices of *16p11.2del* +/+ and *16p11.2del* +/- embryos transfected with a combination of TAOK2β and tDimer2 plasmids via IUE at E14.5-E15 were isolated two days later. The transfected cortical regions were identified by the red fluorescent signal and dissected on a cold surface under a stereomicroscope (Olympus SZX16) connected to a UV light source. Afterwards, the cells were prepared and cultured for 7DIV.

### Polysome profiling

Frozen dissected cortices (prefrontal and somatosensory) from both genders of four weeks old *Taok2* +/+, *Taok2* +/-, and *Taok2* -/- mice or *16p11.2del* +/+, *16p11.2del* +/- and *delTgTaoK2* mice were used for profiling. Polysome profiling was conducted as previously described ^60^ and 10 mM MgCl_2_ was used in preparation of the sucrose gradients. To confirm the association of TAOK2β with polysomes, we followed the standard procedures used for polysome profiling of the cortex, and EDTA was added directly to the pre-cleared cytoplasmic lysates (final concentration 30 mM) which were maintained on ice for 10 minutes before loading into the sucrose gradient prior to centrifugation.

48 hours post-transfected and control N2a cells with 70-80% confluency at the harvesting time were treated with 100 µg/ml CHX at 37°C for 3 minutes before lysis. Cells were washed twice with ice-cold 1x phosphate buffer saline (PBS) containing 100 µg/ml CHX. The cells were gently scraped from the plate in the residual PBS/CHX with a cell scraper. The cells from two 10 cm plates for each gradient were pooled together and spun down at 1,000 x g for 5 minutes at 4°C. A cold lysis buffer containing 20 mM Tris HCl (pH 7.4), 100 mM KCl, 5 mM MgCl_2,_ 0,5 mM DTT, CHX 100 µg/mL, 40 U/mL RNAsin, 20 U/mL Superasin, 25 U/ml Turbo DNase I (Invitrogen), 1% NP-40, 1% Triton-X 100, 1x cOmplete protease inhibitor cocktail (Roche) and phosSTOP (Roche) was added to the pelleted cells and incubated on ice for 10 minutes. Afterwards, cells were triturated 10 times using a 27-gauge syringe, and the lysates were cleared by centrifugation at 2,000 x g for 3 minutes, followed by 16,900 x g for 7 minutes. The cleared cytoplasmic supernatants were transferred into an ice-cold microcentrifuges tube, the OD_260_ of each lysate was measured, and equal OD unit volumes were loaded into 17.5% − 50 % sucrose gradients prepared in a buffer containing 20 mM Tris–HCl (pH 7.4), NaCl 100 mM, 5 mM MgCl2, 1 mM DTT and CHX 100 µg/mL. Sucrose gradients preparation, sample ultracentrifugation, gradient profiling, and collections of the fractions were done as previously described ^60^.

1.5-2.0 × 10^9^ LCLs per gradient were spun down by centrifugation at 1,000 rpm at RT and then treated with a prewarmed medium supplemented with 100 µg/ml CHX at 37°C for 5 minutes. After CHX treatment, the cells were pelleted by centrifugation at room temperature (RT) and washed 2 times with ice-cold 1x PBS supplemented with 100 µg/ml CHX. The pelleted cells were lysed in the same polysome lysis buffer used for lysis N2a cells containing 10 mM MgCl2 and profiled under similar conditions. Polysomes to monosome (P/M) ratio was calculated as described previously ^60^. The P/M ratios generated from profiles of a certain experiment were normalized to the mean P/M ratio of the control profiles within the same experiment followed by statistical analysis for all pooled replicates.

### Measurement of protein synthesis by SUnSET assay

#### Puromycin treatment

To measure the protein synthesis in the acute brain slices, we followed a previously described protocol ^15, 16^. Briefly, 400 μm coronal brain slices were prepared from four weeks old *Taok2* +/+, *Taok2* +/-, and *Taok2* -/- mice with a vibratome in a cutting solution containing 85 mM NaCl, 75 mM sucrose, 2.5 mM KCl, 25 mM glucose, 1.25 mM NaH_2_PO_4_, 4 mM MgCl_2_, 0.5 mM CaCl_2_, 24 mM NaHCO_3_ at low temperature. Slices were recovered at 32 °C in an incubation chamber with the cutting solution for 30 minutes followed by another 30 minutes with perfused oxygenated artificial cerebrospinal fluid (ACSF) containing 127 mM NaCl, 25 mM NaHCO_3_, 25 mM D-Glucose, 1.25 mm NaH_2_PO_4_, 2.5 mM KCl, 1 mM MgCl_2_, and 2 mM CaCl_2_ at pH 7.4, before the treatment. Slices were then treated with puromycin (10 µg/ml) for 90 minutes to label the newly synthesized proteins. The incubation media with puromycin was removed followed by 2 successive washes with oxygenated ACSF for 10 minutes each before lysis for western blot processing or fixation for immunohistochemical investigations. To verify the detected signal of the newly synthesized protein by anti-puromycin antibody and whether it is protein synthesis-dependent, slices from wild-type animals were pre-treated with 100 μm/ml CHX (a translation inhibitor) for 30 minutes before puromycin treatment.

Primary cortical neurons (7DIV or 21DIV), N2a cells (control cells or cells transfected with TAOK2β or TAOK2βA135P plasmids) and 5 x 10^6^ LCLs were treated with puromycin (10 µg/ml) in a pre-warmed medium and incubated for 10 minutes at 37°C, 5% CO2. To verify the detected signal of the newly synthesized protein by anti-puromycin antibody, cells were pre-treated with 50µg/ml of CHX for 5 minutes at 37°C, 5% CO2, and then followed by puromycin treatment. After puromycin treatment, cells were rinsed quickly with a standard medium. Afterwards, cells were either fixed for immunostaining or washed twice with ice-cold 1x PBS, pelleted by centrifugation at 1,000 r.p.m for 5 minutes at 4°C and processed with western blotting for SUnSET analysis.

#### Immunofluorescence SUnSET analysis

Acute brain slices were fixed with 4% paraformaldehyde in PBS (4% PFA) for 1.5 hours at RT. Slices were permeabilized with 0.3% Triton X-100 in PBS, followed by incubation in blocking buffer (5% Donkey Serum in TBS, 0.5 % Triton) for 1.5 hours at RT. Primary antibody incubations were conducted for 48 hours at RT with gentle shaking. After 3 washes with 1x PBS, slices were incubated with anti-Alexa Fluor conjugated secondary antibodies in 1 x PBS with 0.3% Triton X-100 for 2 hours at RT. Slices were washed three times in 1x PBS and incubated with DAPI for 30 minutes at RT. Slices were washed 3 times with 1x PBS and then mounted using a mixture of Mowiol and DABCO. Neurons or N2a cells were fixed with 4% PFA in PBS containing 4% sucrose at RT for 10 minutes at RT followed by 3 washes with 1x PBS. Cells were permeabilized with 1x PBS containing 0.3% Triton X-100 and incubated in blocking buffer (5% Donkey Serum in TBS, 0.3 % Triton) for 1.5 hours at RT. Primary antibody incubations were carried out overnight at 4°C and secondary antibodies were added for two hours at RT. After washing, coverslips were stained with DAPI and mounted on glass microscope slides using Fluoromount-G (Southern Biotech).

Slices were imaged by Zeiss LSM 900 confocal laser microscope equipped with Plan-Apochromat 20x/0.8 objective. Z-series images were acquired with a step size of 1μm in-between z sections. Cells were imaged by a Nikon microscope (Eclipse, Ti) using an inverted 60 X oil immersion objective (NA 1.4). Cells were identified by their morphology or immunolabelling by anti-beta tubulin III, anti-TAOK2β or anti-Myc-tag antibodies or fluorescent markers (tDimer or EGFP). Z-series images were acquired with a step size of 0.3 μm in-between z sections. Image acquisition settings were maintained similarly within the experiment. The brightness and contrast of the images were adjusted with the same setting using Photoshop CS 8.0 (Adobe systems) or ImageJ.

All z sections for the entire cell were summed together with the maximum intensity using the z project tool in the ImageJ. Afterwards, mean puromycin fluorescent intensity in the cell body was measured in randomly selected labelled cells by ImageJ. Only individual cells with no adjacent cells were analyzed to avoid false-positive signals from the other cells. Transfected neurons expressing tDimer and TAOK2β, and the nearby non-transfected neurons from one coverslip were selected and processed in parallel to control the conditions. Only N2a cells expressing Myc-tag plus GFP signals were analyzed.

#### Western blot SUnSET analysis

Micro-dissected cortices from the acute slices or N2a cells growing on a 10 cm culture dish or 5 x 10^6^ LCLs following the puromycin treatment were lysed in sterile-filtered RIPA buffer (50 mM Tris-HCl (pH 7.4), 150 mM NaCl, 1mM EGTA, 1% NP-40, 0.25% Sodium deoxycholate) supplemented with 1x cOmplete protease inhibitor cocktail (Roche) by sonication 5x every 10 seconds. The lysates were cleared by centrifugation at 20.000 x g for 10 minutes at 4°C. Equal amounts of protein were resolved on 10% sodium dodecyl sulphate (SDS) polyacrylamide gel electrophoresis (PAGE) and immunoblotted against puromycin and β-actin. Puromycin protein signal in the whole lane was normalized against β-actin to calculate the puromycin/β-actin ratio using ImageJ. Revert™ 700 Total Protein Stain for Western Blot Normalization (Licor) was used to show the equal protein loading.

### Immunoprecipitation (IP) using TAOK2β antibody

The cortex of *Taok2* +/+ mouse or N2a cells 48 hours post transfection with wild-type TAOK2β were lysed in ice-cold lysis RIPA buffer (pH 8.2) containing 1x cOmplete protease inhibitor cocktail (Roche). The cellular homogenates were cleared from the cell debris by centrifugation at 13,000 rpm for 10 minutes at 4°C. 1000 µg cleared homogenate was incubated with polyclonal rabbit anti-Taok2β or polyclonal rabbit anti-IgG antibodies prebound to Dynabeads Protein A (Invitrogen) undergoing rotation (25 rpm) overnight at 4°C. The supernatant was removed using a magnetic holder and the immunoprecipitated complexes were washed 3 times with a pre-cooled lysis buffer at 4°C. Following another 3 washes with Tris-HCl pH 8.2 at 4°C, the immunoprecipitated complex was analyzed by LC-MS/MS. For analysis of the immunoprecipitated complex from the cytoplasmic lysates of N2a cells by SDS-PAGE, 50 µl Laemmli sample buffer was added to the immunoprecipitated complex and boiled for 10 minutes at 95°C in a heat block to elute the proteins. Afterwards, the eluted proteins were loaded on 7.5-20% SDS-polyacrylamide gel and processed with the routine WB analysis and immunodetection using anti-Taok2β and anti-eEF2 antibodies.

### Neuronal proximity-based proteomic system

to identify protein-protein interaction (PPI) in DIV18 neuron, we followed our previously described protocol ^12^.

### Trichloroacetic acid (TCA) protein extraction from polysomes fractions for western blot analysis

Every 2-3 sucrose fractions were pooled together as indicated in the scheme (Fig. 2a, b). An equal volume of 20% TCA was added to the sample in a microcentrifuge tube and incubated for 30 minutes on ice. Afterwards, the tubes were centrifugated at 14,000 rpm for 5 minutes. The supernatant was gently removed, and the pellet was washed with 500 µl cold acetone and spun down at 14,000 rpm for 5 minutes. After two-acetone washes, the pellets were dried by placing the tubes in a 50°C heating block for 5 minutes. All procedures were performed under a safety hood. For SDS-PAGE, 50 µl 2x Laemmli sample buffer was added to each pellet. The pellets of the first 3 pooled fractions were dissolved in 150 µl 2x Laemmli sample buffer. The sample was boiled for 10 minutes at 95°C in a heat block and 30 µl were loaded onto 10% SDS-PAGE gel for routine western blotting

### In vitro kinase assay

Ascending concentrations of 25, 50, 100 and 150 ng of recombinant human protein TAOK2 (aa 1-314) with N-terminal GST tag (Signal-Chem, T25-11G-10) were incubated with 300 ng of recombinant human protein eEF2 with N-terminal 6xHis-tag (MyBioSource, MBS1213669) in a 25 µl kinase assay buffer I (Signal-Chem, K01-09) supplemented with cOmplete protease inhibitor cocktail (Roche) and phosSTOP (Roche). For control, 300 ng of the recombinant human protein eEF2 protein was incubated in the reaction buffer without adding the recombinant human protein TAOK2 or 150 ng of TAOK2 protein without the eEF2 protein. The kinase reaction was initiated by adding ATP to a final concentration of 350 µM and incubation at 30°C for 45 minutes. In the control reaction, 150 ng of TAOK2 was used under the same reaction conditions without the substrate. The reaction was terminated by adding Laemmli sample buffer and subjected to western blot analysis to detect the phosphorylation of eEF2 at its Thr56 residue by using an anti-phospho-Thr56 specific rabbit antibody.

### Immunoblot

30 µl sample with Laemmli buffer after TCA protein precipitation from each polysomes fraction or 30-50 µg protein determined by Pierce™ BCA Protein Assay Kit according to the manufacturer’s guidelines from cellular or brain lysates were subjected to standard SDS-PAGE. Immunoblotting to PVDF membrane was performed with the standard wet transfer method. Blots were probed with primary antibodies following the antibody’s datasheet instructions. The signals were visualized by HRP conjugated anti-IgG secondary antibodies, imaged by enhanced chemiluminescence NTAS, ChemoStar, ECL Imager and analyzed using software from Fiji Software. Proteins of interest were normalized to the β-actin signal and the phospho-specific signals were normalized to their respective total protein intensity after membrane stripping and redevelopment.

### Protein purification from polysomes to detect Taok2 isoforms by proteome analysis

Sucrose polysome fractions 11-22 (that represent light and heavy polysomes in the polysome profile) from *Taok2* +/+ mouse cortex were collected and pooled together. The proteins were purified from the sucrose and concentrated up to 200 µl final volume by several washes with ddH_2_O using Amicon Ultra −15 centrifugation filter (Millipore) at 4°C. Protein concentration in the sample was determined and equal amounts of protein were loaded into 10% SDS-PAGE gel. Following electrophoresis, the protein bands were visualized by Coomassie (Roti®-Blue) staining solution (Roth) for 30 minutes at RT. The gel was washed several times with 25% methanol solution until the bands were visible. The bands at approximate 100-160 kDa corresponding to Taok2 molecular weight were excised, digested, and analyzed by LC-MS/MS.

### Proteome analysis for the purified protein from polysomes and the immunoprecipitated proteins with Liquid chromatography-mass spectrometry (LC-MS/MS)

Tryptic in-gel digestion was conducted following a previously described protocol ^61^. Shrinking and swelling of gel pieces were performed with 100% acetonitrile (ACN) and 100 mM ammonium bicarbonate (NH_4_HCO_3_). The in-gel reduction was conducted with 10 mM dithiothreitol (DTT) (dissolved in 100 mM NH_4_HCO_3_). Alkylation was performed at room temperature with 55 mM iodoacetamide (dissolved in 100 mMNH_4_HCO_3_). Digestion of proteins in the gel pieces was done by covering them with a trypsin solution containing (8 ng/µL sequencing-grade trypsin (Promega), dissolved in 50 mM NH_4_HCO_3_with 10% ACN). The mixture was incubated overnight at 37°C. Tryptic peptide products were further yielded by extraction with 2% formic acid (FA) and 80% ACN. The extract was evaporated. For LC-MS/MS analysis, samples were resuspended in 20 µL of 0.1 % FA. Protein digestion of the immunoprecipitated samples was performed on the magnetic beads. Therefore, 1% sodium deoxycholate (SDC) in 100 mM triethylammonium bicarbonate was added and the samples were boiled at 95°C for 5 minutes. DTT was added to the sample (final concentration 10 mM) and incubated at 60°C for 30 minutes. Iodoacetamide was added to the samples (final concentration 20 mM) and incubated at 37°C for 30 minutes. Proteins were digested with trypsin (sequencing grade, Promega) at 37°C overnight. After digestion, samples were placed on a magnetic rack, left for 1 minute to settle and the supernatant was transferred to a new tube, 1% FA was added to stop digestion and precipitate SDC. Samples were then centrifuged at 16,000 x g for 5 minutes, and the supernatant was transferred to a new tube and dried in a vacuum centrifuge.

Samples were reconstituted in 0.1% FA and transferred into a full recovery autosampler vial (Waters). Chromatographic separation was achieved on a Dionex Ultimate 3000 UPLC system (Thermo Fisher Scientific) with a two-buffer system (buffer A: 0.1% FA in water, buffer B: 0.1% FA in ACN). Attached to the UPLC was an Acclaim PepMap 100 C18 trap (100 µm x 2 cm, 100 Å pore size, 5 µm particle size, Thermo Fisher Scientific) for desalting a purification followed by a nanoEase M/Z peptide BEH130 C18 column (75 µm x 25 cm, 130 Å pore size, 1.7 µm particle size, Waters). Peptides were separated using a 60 min gradient with increasing ACN concentration from 2% − 30% ACN. The eluted peptides were analyzed on a quadrupole orbitrap Ion trap tribrid mass spectrometer (Fusion, Thermo Fisher Scientific) in data-dependent acquisition (DDA). The fusion was operated at top speed mode analyzing the most intense ions per precursor scan (2×10^5^ ions, 120,000 Resolution, 120 ms fill time) within 3 s and were analyzed by MS/MS in the ion trap (HCD at 30 normalized collision energy, 1×10^4^ ions, 60 ms fill time) in a range of 400 – 1300 m/z. A dynamic precursor exclusion with a speed mode of 20 s was used.

The acquired DDA LC-MS/MS data were searched against the Uniprot mouse protein database (release October 2020, 17,053 protein entries) and the TAOK2 splice variant 2 (Q6ZQ29-2) using the Sequest algorithm integrated into the Proteome Discoverer software version 2.4 in label-free quantification mode. The match between runs was enabled and performing chromatographic retention re-calibration for precursors with a 5 min retention time tolerance, no scaling, and no normalization for extracted peptide areas was done. The following parameters were applied in the searches: Mass tolerances for precursors were set to 10 ppm and fragment mass tolerance was 0.6 DA. Carbamidomethylation was set as a fixed modification for cysteine residues and the oxidation of methionine, pyro-glutamate formation at glutamine residues at the peptide N-terminus as well as acetylation of the protein N-terminus, loss of methionine at the protein N-terminus and the acetylation after methionine loss at the protein N-terminus were permitted as variable modifications for the search. Peptide areas were summed to protein areas and used for quantitative analysis. Only the peptides with high confidence (false discovery rate < 1% using a decoy database approach) were accepted as identified. Peptide areas were summed to protein areas and used for quantitative analysis. Protein areas were imported into Perseus software version 1.5.8 for statistical analysis.

N2a cells and mouse cortices were run in duplicates with their corresponding IgG controls. The resulting protein abundance was first averaged between the respective samples, and only proteins with a non-zero count in at least one of the duplicates in the IgG control were kept for further analysis in R with customs scripts (v. 4.1.0, available upon request). The filtered protein abundances comprised 293 proteins for N2a cells and 194 proteins for the cortex. In the parallel PPI analysis from cultured neurons, we performed a t-test for further confidence of protein detection with a p-value of 0.05 and a positive fold change compared to the luciferase control as cut-off, resulting in identified 150 proteins in neuronal cell culture for 18DIV. The ORA depiction as cnetplot between the three MS approaches (N2a cells overexpressing TaoK2β, cortex, DIV18 neurons) was generated using enrichplot R package v. 1.12.3. The dotplot was generated by applying compare Cluster function from R package clusterProfiler v. 4.0.5 with a p-value cutoff of 1 and enrich GO parameter.

### Total and phosphoproteomics of *Taok2* -/- mouse brain regions

Frozen tissue was punched from 1mm-slices from the prefrontal cortex, ventral striatum, and hippocampus of P21 *Taok2* +/+ (6), *Taok2* +/- (5) or *Taok2* -/- (5) mouse brain. Tissue was lysed for 30 min on ice in lysis buffer containing 8M urea, 50 mM HEPES, pH 8.0, 1 mM DTT, cOmplete protease inhibitor cocktail (Roche) and phosSTOP (Roche) followed by 3x 10s sonication. 100 µg of protein was reduced with 5 mM DTT for 30 min at 37°C and alkylated with 15 mM iodoacetamide for 30 min at RT in the dark. Proteins were digested with 4 ug Trypsin/Lys-C mix (Promega V5073) for 3-4 hrs at 37°C, diluted 4x with 50 mM HEPES pH 8.0 to reactivate trypsin and incubated o/n at 37°C. Samples were desalted using Pierce Peptide Desalting Columns (Thermo Scientific 89851) and vacuum dried completely. Dried peptides were dissolved in HEPES, pH 8.5 and labelled with 0.5 mg TMTpro 16plex Isobaric Label Reagent Set (Thermo Scientific A44522). Individually labelled samples were combined and desalted using a Pierce Peptide Desalting Column (Thermo Scientific 89851). Labelled peptides were dried and used for Sequential Enrichment with Metal Oxide Affinity Chromatography (SMOAC). First, phosphopeptides were enriched using High-Select TiO2 Phosphopeptide Enrichment Kit (Thermo Scientific A32993). The flowthrough was subsequently enriched using the High-Select Fe-NTA Phosphopeptide Enrichment Kit (Thermo Scientific A32992). Elutions of both steps were combined and used for phosphoproteomics mass spectrometry. The final flowthrough was used for total proteomics mass spectrometry.

50µg dried peptides were redissolved in 100 µl solvent A (10mM ammonium bicarbonate in water/ACN (98:2, v/v), pH 5.5). Fractionation was performed by RP-HPLC (Agilent series 1200) connected to a Probot fractionator (LC Packings), where 95 µl was loaded onto a 4 cm pre-column (made in-house, 250 µm internal diameter (ID), 5 µm C18 beads, Dr. Maisch) for 10 minutes followed by separation on a 15 cm analytical column (made in-house, 250 µm ID, 3 µm C18 beads, Dr Maisch) with a linear gradient from 0% solvent B (10mM ammonium bicarbonate in water/ACN (30:70, v/v), pH 5.5) up to 100% solvent B in 120 minutes at a flow rate of 3 µl/min. 1-minute fractions were pooled every 12 minutes into 12 pooled fractions, vacuum dried and stored at −20°C until LC-MS/MS analysis.

Each fraction was solubilized in 30 µL loading solvent A (0.1% TFA in water: ACN (98:2, v:v)) moments before analysis and 12/13 µl was injected for LC-MS/MS analysis on an Ultimate 3000 RSLCnano system in-line connected to an Orbitrap Fusion Lumos mass spectrometer (Thermo). Trapping was performed at 10 μl/min for 4 min in loading solvent A on a 20 mm trapping column (made in-house, 100 μm internal diameter (I.D.), 5 μm beads, C18 Reprosil-HD, Dr. Maisch, Germany). The peptides were separated on a 200 cm µPAC™ column (C18-endcapped functionality, 300 µm wide channels, 5 µm porous-shell pillars, inter pillar distance of 2.5 µm and a depth of 20 µm; Pharmafluidics, Belgium). It was kept at a constant temperature of 50°C. Peptides were eluted by a linear gradient reaching 55% MS solvent B (0.1% FA in water/acetonitrile (2:8, v/v)) after 85 min and 99% MS solvent B at 90 min, followed by a 10-minute wash at 99% MS solvent B and re-equilibration with MS solvent A (0.1% FA in water). For the first 15 min the flow rate was set to 750 nl/min after which it was kept constant at 300 nl/min.

Total proteomics was performed using an SPS-MS3 method with the mass spectrometer operated in data-dependent mode with a top speed of three seconds. Full-scan MS spectra (375-1500 m/z) were acquired at a resolution of 120,000 in the Orbitrap analyzer after accumulation to a target AGC value of 400,000 with a maximum injection time of 50 ms. The precursor ions were filtered for charge states (2-7 required), dynamic exclusion (60s; +/- 10 ppm window) and intensity (minimal intensity of 5E4). The precursor ions were selected in the quadrupole with an isolation window of 0.7 Da and accumulated to an AGC target of 1E4 or a maximum injection time of 50 ms and activated using CID fragmentation (35% NCE). The fragments were analyzed in the Ion Trap Analyzer at a turbo scan rate. 10 most intense MS2 fragments were selected in the quadrupole using MS3 multi-notch isolation windows of 3 m/z. An orbitrap resolution of 60,000 was used with an AGC target of 1E5 or a maximum injection time of 118 ms and activated using HCD fragmentation (65% NCE).

Phosphoproteomics was performed like the total proteomics samples, but instead using an SPS-MS3 with a multistage activation method. The fragments were analyzed in the Orbitrap Analyzer with a resolution of 30,000. 10 most intense MS2 fragments were selected in the quadrupole using MS3 multi-notch isolation windows of 3 m/z. An orbitrap resolution of 60,000 was used with an AGC target of 1E5 or a maximum injection time of 105 ms and activated using HCD fragmentation (65% NCE).

Database search was done with the MaxQuant software (v 1.6.2.6) using the Andromeda search engine with the default search settings including an FDR set at 1% on both the peptide and protein level. Reporter ion MS3 was set as search type and TMT16pro isobaric labels were manually entered with appropriate correction factors. Reporter mass tolerance was set at 0.003 Da. Spectra were searched against the Mus musculus database of UniProt (August 2020) and the common contaminant list provided by the software. The MS1 mass tolerance for the first search and the main search was set to 20 ppm and 4.5 ppm respectively. The MS2 match tolerance is set at 20 ppm. Enzyme specificity was set to Trypsin/P with digestion at the C-terminal of Arg and Lys residues, even when they were followed by a Pro residue, with a maximum of two missed cleavages. Variable modifications were set to oxidation (Met) and protein N-terminal acetylation. Carbamidomethylation of Cys-residues was set as a fixed modification. For phosphoproteomics, phosphorylation at serine, threonine or tyrosine residues was set as a variable modification. A minimum of one peptide (razor or unique) was required for identification with a minimum length of 7 residues. Second peptide search was allowed as well as the matching between runs option (from and to) using a 0.7-minute match time window and twenty-minute alignment window.

Data were further processed in Perseus v.1.6.15.0. Protein Groups and Phospho (STY) sites from the MaxQuant search were loaded and site tables expanded. Potential contaminants, reversed hits and proteins that were identified only by site, were removed. Reporter intensities were log2 transformed and filtered for rows containing only valid values. Columns were normalized by subtracting the median. Potential TAOK2 substrates were annotated using the pT-X-X-[RKH] motif. A multi-scatter plot and principal component analysis were calculated to determine the quality of the runs. A two-sample t-test was performed comparing the WT and KO conditions with a permutation-based FDR of 0.05 and an S0 value of 0.1 for truncation with a total of 250 randomizations.

Gene set enrichment analysis (GSEA) was performed on the total proteomics dataset ranked by signed log-transformed P values using the Molecular Signatures Database GO Biological Process collection (MsigDB, v7.5.1). The pre-ranked list was given as input to GSEAPreranked (v4.2.2) with default parameters. Network analysis was done with EnrichmentMap and AutoAnnotate in Cytoscape (v3.9.1) using an FDR cutoff of 0.1 and similarity cutoff of 0.375.

## Statistical analysis

The GraphPad Prism (version 9) analytic software was used to perform all the statistical analyses. Several biological replicates, experiments and statistical tests used for the comparison were mentioned within the figure legends.

**Supplementary Table 1:** TAOK2β interacting proteins (IP-MS analysis).

**Supplementary Table 2:** TAOK2 isoforms in polysomes of wild-type mouse cortex.

**Supplementary Table 3:** TAOK2 total proteomics.

**Supplementary Table 4: TAOK2 Phosphoproteomics.** Supplementary Tables are posted online as Excel files.

---

**Supplementary Figure 1:**
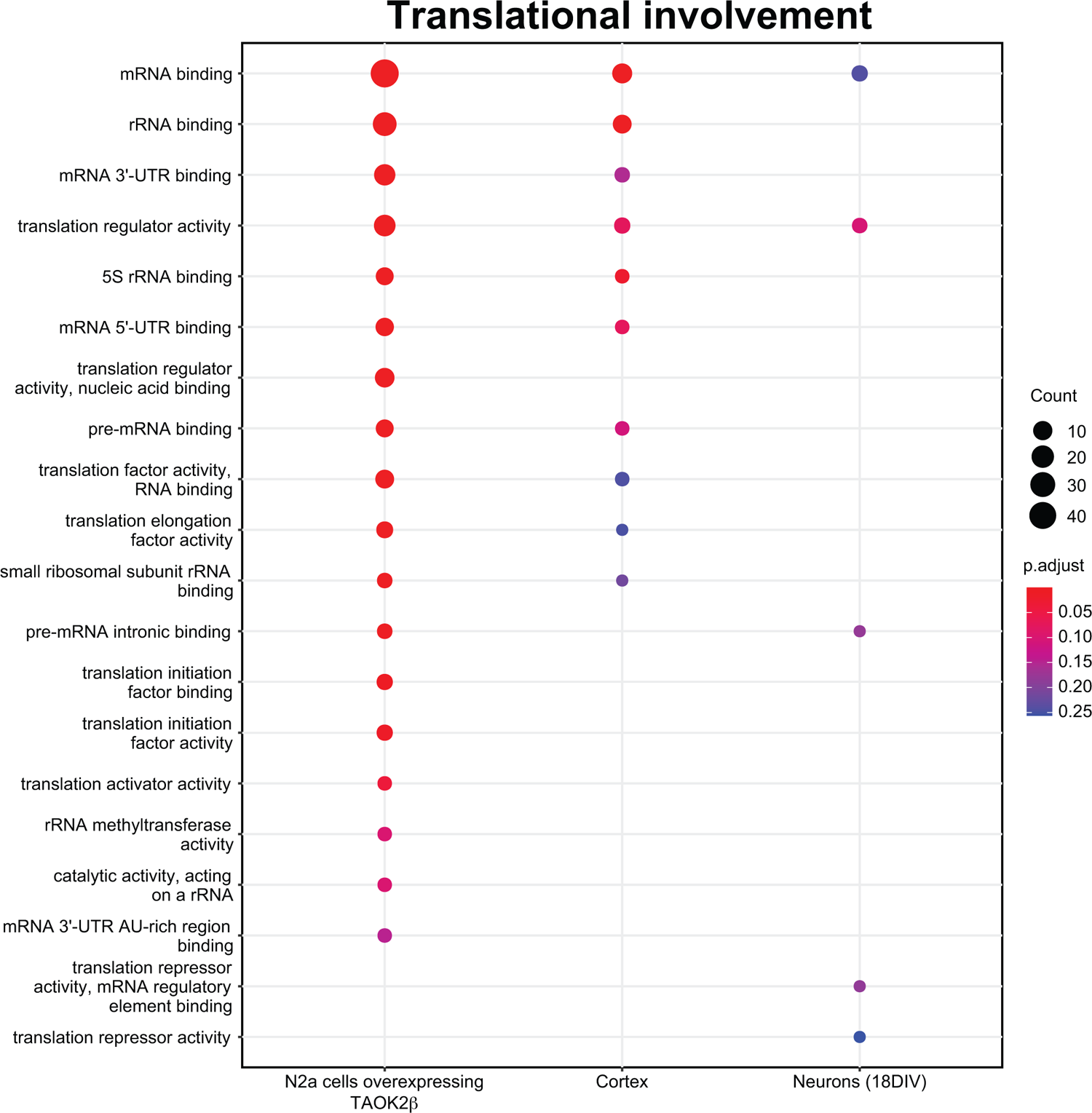
TAOK2β associates with different processes of translation regulation in N2a cells expressing TAOK2β, mouse cortex and cortical neurons. A detailed overview of the translational involvement of TAOK2β interacting proteins indicates the contribution of TAOK2β to different translational processes such as RNA binding, translation initiation and translation elongation.

**Supplementary Figure 2:**
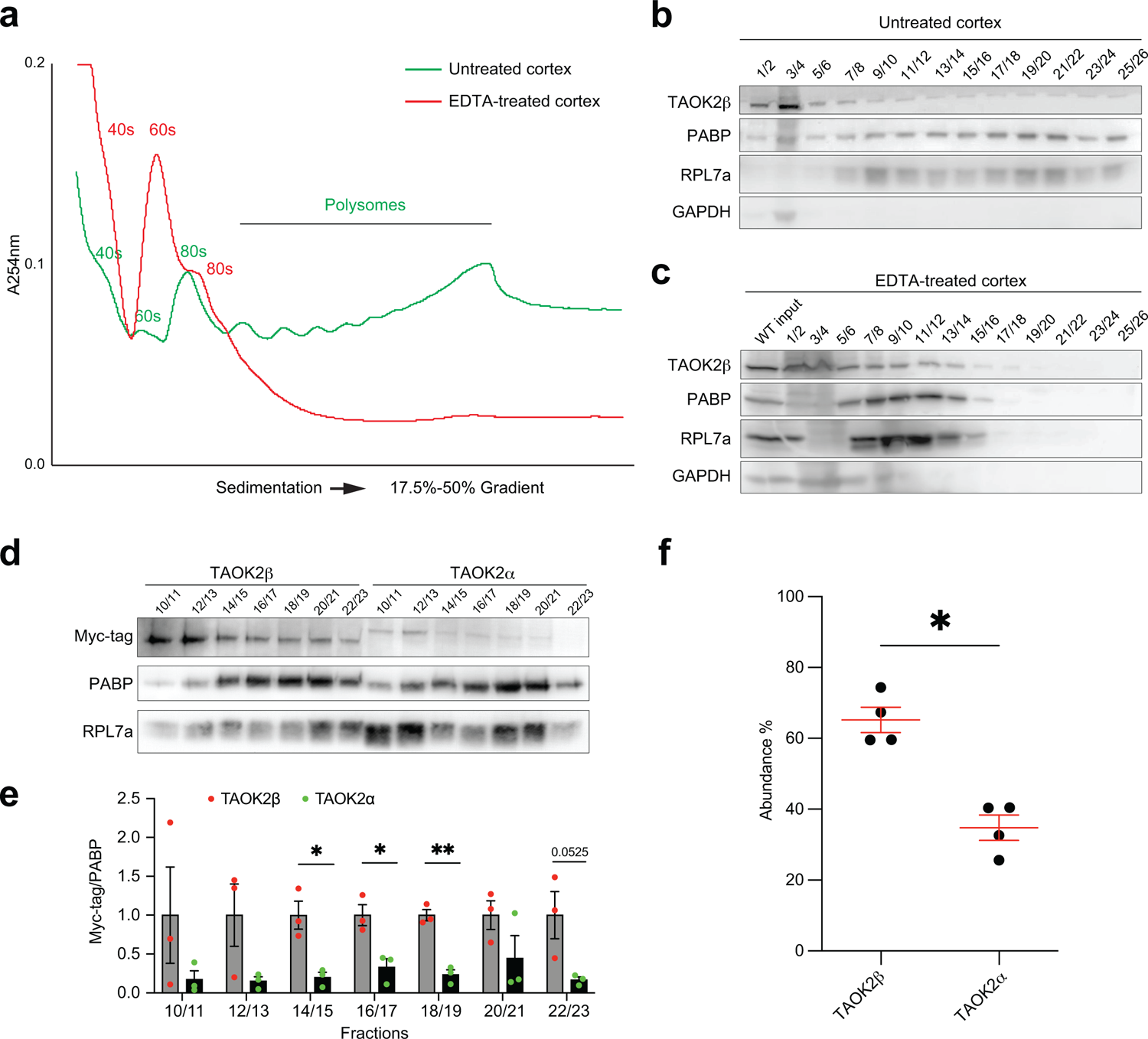
TAOK2β associates with the actively translating polyribosome complex. **(a)** Polysome profiles of *Taok2* +/+ cortices under normal basal conditions (green) or after EDTA treatment (red) to disrupt polysomes. EDTA treatment causes dissociation of ribosomes leading to the disappearance of polysomes and an increase in peaks of monosomes, and the ribosomal subunits 40S and 60S. **(b, c)** Immunoblots of cortical polysome fractions pooled as indicated (Figure 2b), showing the presence of TAOK2β across all fractions from polysomes in the untreated cortex. Note, the presence of TAOK2β coincides with PAPB and RPL7a expression (positive controls) across all heavier polysome fractions from the untreated cortex. Following EDTA treatment, the distribution of PABP1, RPL7a, and TAOK2β signals are no longer detectable in the heavier polysomes fractions and instead accumulate in the unbound and monosome fractions. GAPDH was used as a negative control. **(d)** Immunoblots for the pooled gradient polysomes fractions (10-23) of N2a cells either transfected with wild-type TAOK2β or TAOK2α plasmids show higher abundance of TAOK2β in polysomes than TAOK2α as detected by anti-Myc-tag antibody. PABP and RPl7a were used as positive controls. **(e)** Quantifications of the immunoblot (d) show a significant increase of densiometric anti-Myc-tag antibody signal normalized to PABP in polysomes of N2a cells expressing wild-type TAOK2β compared to cells expressing TAOK2α. Number of biological replicates: n = 3 per condition from different independent transfection experiments; *P < 0.05, **P < 0.01, SEM. error bars, unpaired t-test. **(f)** Quantification of the Taok2 isoforms in polysomes isolated from *Taok2* +/+ mouse cortices analyzed by LC-MS/MS showing the increased abundance of the Taok2β (65.21%) compared to the Taok2α isoform (34.79%). n = 4 cortices from 4 weeks old *Taok2* +/+, *P < 0.05, SEM. error bars, unpaired t-test.

**Supplementary Figure 3:**
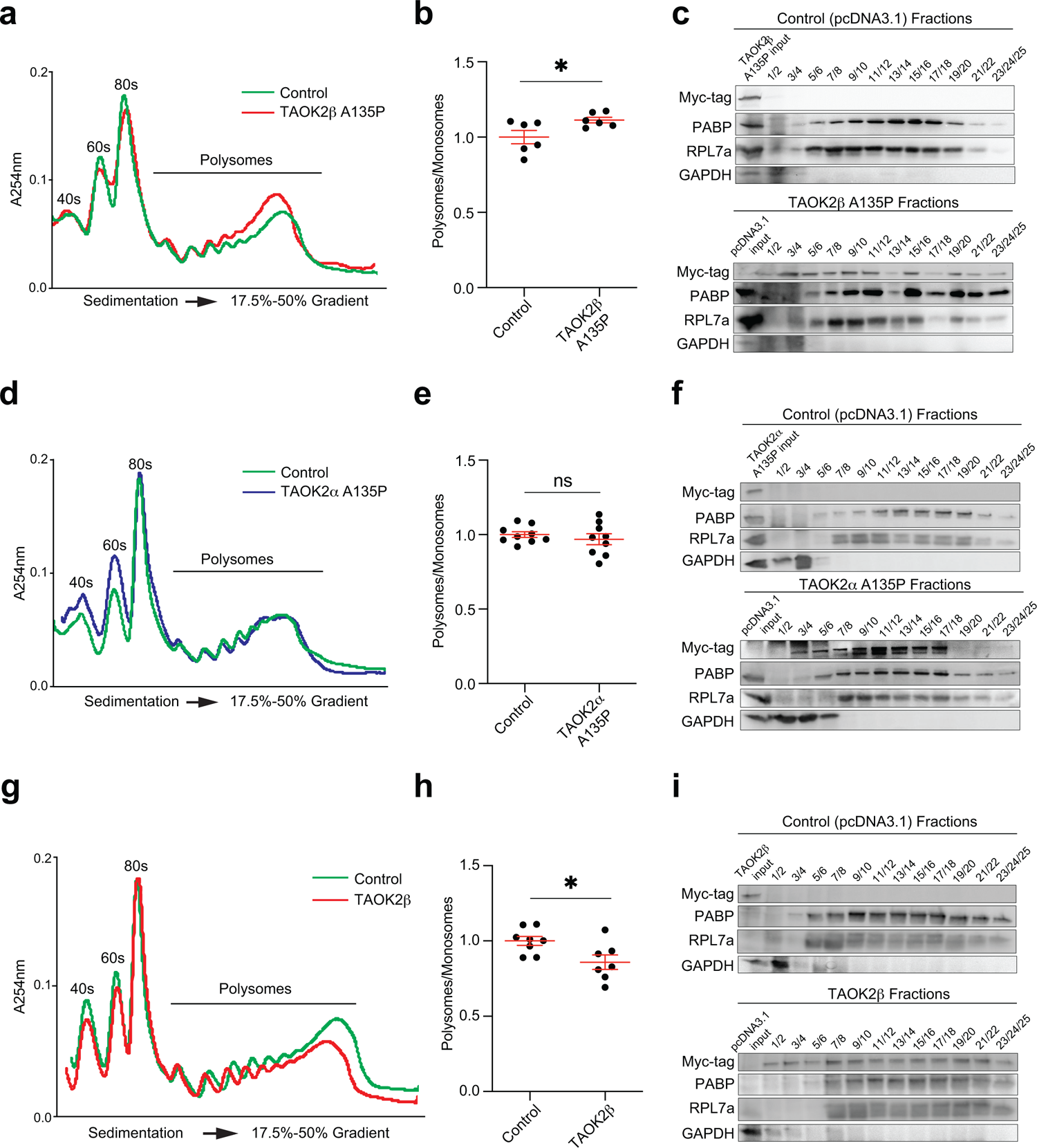
General translation is enhanced in cells expressing TAOK2β A135P not TAOK2α A135P and TAOK2β represses general translation. **(a)** Overlay of polysome profiles of N2a cells transfected with TAOK2βA135P or control plasmids shows an increase in the P/M ratio of the profile from cells expressing TAOK2βA135P **(b)** Quantifications of the normalized P/M ratios of polysome profiles reveal a statistically significant increase in the P/M ratios for profiles of cells expressing TAOK2βA135P. Number of biological replicates, TAOK2βA135P transfected cells n = 5, control n = 5; SEM. error bars; *P < 0.05, unpaired t-test. **(c)** Immunoblots of the pooled gradient polysomes fractions of N2a cells either transfected with TAOK2βA135P or control plasmids show the presence Myc-tag in polysomes of cells expressing TAOK2βA135P (lower). PABP1 and RPl7a were used as positive controls and GAPDH as a negative control. Input from the total cytoplasmic lysate, either from cells expressing TAOK2βA135P or control cells expressing pcDNA3.1-myc-tag, was used to verify the specificity of the antibody and the transfection efficiency. **(d)** Overlay of polysome profiles of N2a cells transfected with TAOK2αA135P or control plasmids. **(e)** Quantifications of the normalized P/M ratios of polysome profiles reveal no significant change in the P/M ratios for profiles of the cells expressing TAOK2αA135P transfected cells and control cells. Number of biological replicates, TAOK2αA135P transfected cells n = 9, control n = 9; SEM. error bars; ns, not significant. P= 0.4575, unpaired t-test. **(f)** Immunoblots of the pooled gradient polysomes fractions of N2a cells either transfected with TAOK2αA135P or control plasmid show the presence of Myc-tag in polysomes of cells expressing TAOK2αA135P (lower). PABP1 and RPl7a were used as positive controls and GAPDH as a negative control. Input from the total cytoplasmic lysate, either from cells expressing TAOK2αA135P or control cells expressing pcDNA3.1-myc-tag, was used to verify the specificity of the antibody and the transfection efficiency. **(g)** Overlay of polysome profiles of N2a cells transfected with wild-type TAOK2β or control plasmids shows a decrease in the P/M ratio of the profile from cells expressing wild-type TAOK2β. **(h)** Quantifications of the normalized P/M ratios of polysome profiles reveal a statistically significant decrease in the P/M ratios for profiles of cells expressing wild-type TAOK2β. Number of biological replicates, wild-type TAOK2β transfected cells n= 7, control n= 8, SEM. error bars, *P < 0.05, unpaired t-test. **(i)** Immunoblots of the pooled gradient polysomes fractions of N2a cells either transfected with wild-type TAOK2β or control plasmids show the presence of Myc-tag in polysomes of cells expressing wild-type TAOK2β (lower). PABP1 and RPl7a were used as positive controls and GAPDH as negative. Input from the total cytoplasmic lysate, either from cells TAOK2β or the control cells expressing pcDNA3.1-myc-tag, was loaded to verify the specificity of the antibody and the transfection efficiency.

**Supplementary Figure 4:**
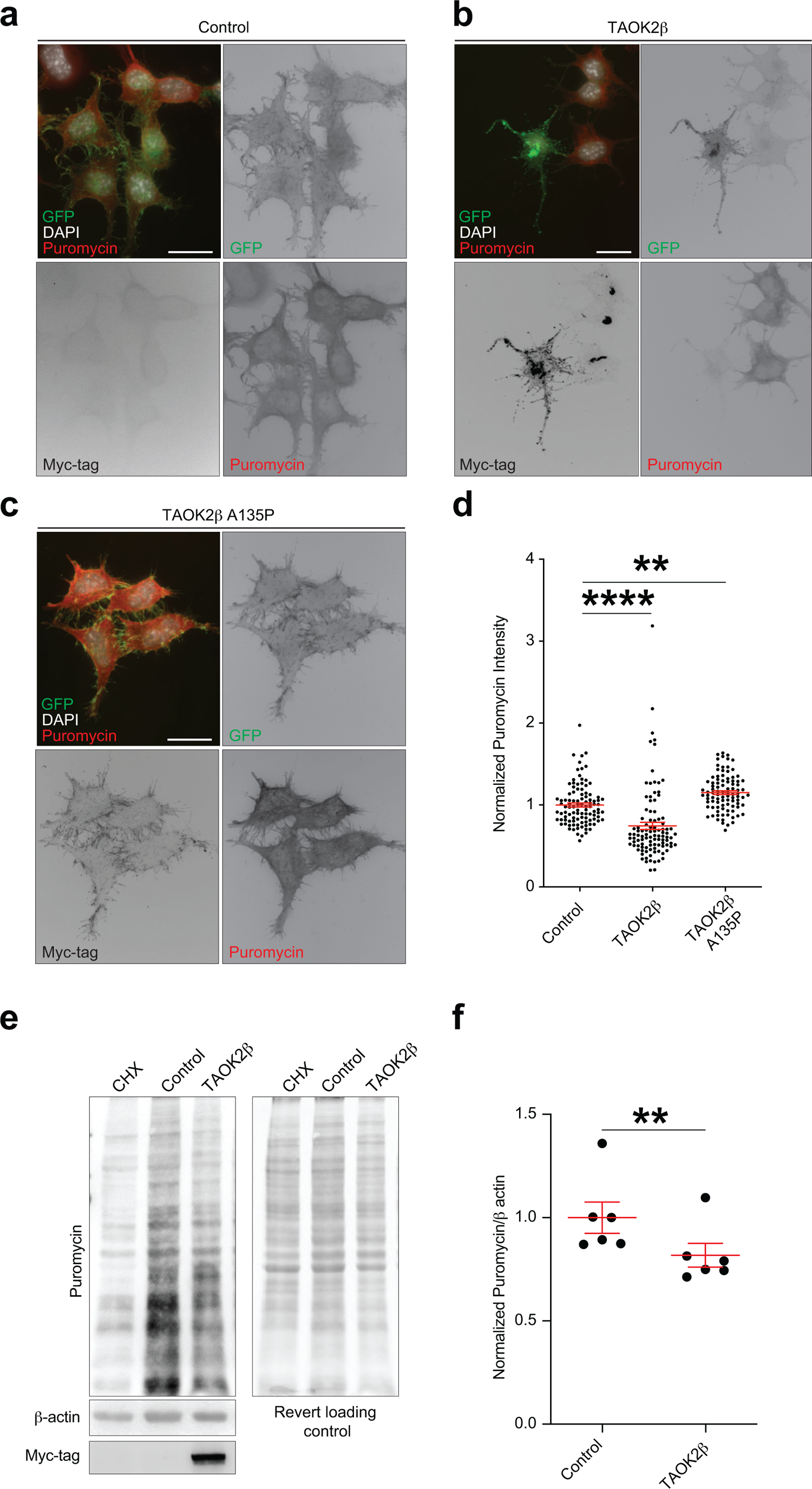
TAOK2β decreases protein synthesis while its A135P mutation increases newly synthesized protein in N2a cells. **(a-c)** Immunostainings of N2a control cells, cells expressing wild-type TAOK2β, and TAOK2βA135P respectively, treated with puromycin to label the newly synthesized protein, fixed and immunostained with anti-puromycin (red) and anti-Myc-tag (grey) antibodies, DAPI (blue) and GFP marker (green). **(d)** Quantifications of the normalized puromycin fluorescence intensity show its decrease in N2a cells expressing wild-type TAOK2β compared to N2a control cells and its increase in cells expressing TAOK2βA135P. Number of cells from 3 independent transfection experiments; control transfected cells n=103, wild-type TAOK2β transfected cells n=102, TAOK2βA135P transfected cells n=93. SEM error bars: ** p < 0.01, **** p < 0.0001, ordinary one-way ANOVA followed by Tukey’s multiple comparisons test. **(e)** Immunoblot of protein lysates from N2a control cells and cells expressing wild-type TAOK2β analyzed by SUnSET assay shows a decreased anti-puromycin antibody signal in cells expressing TAOK2β. Cycloheximide (CHX) was added to non-transfected cells before being treated with puromycin to verify the specificity of the antibody signal. Revert loading control protein stain was used to show equal protein loading. **(f)** Quantifications of the immunoblot SUnSET assay of N2a control cells and cells expressing wild-typeTAOK2β show a significantly decreased densiometric anti-puromycin antibody signal normalized to β-actin in TAOK2β transfected cells. Number of biological replicates n=6 per condition from 3 different independent transfection experiment, SEM error bars; ** p < 0.01, paired t-test.

**Supplementary Figure 5:**
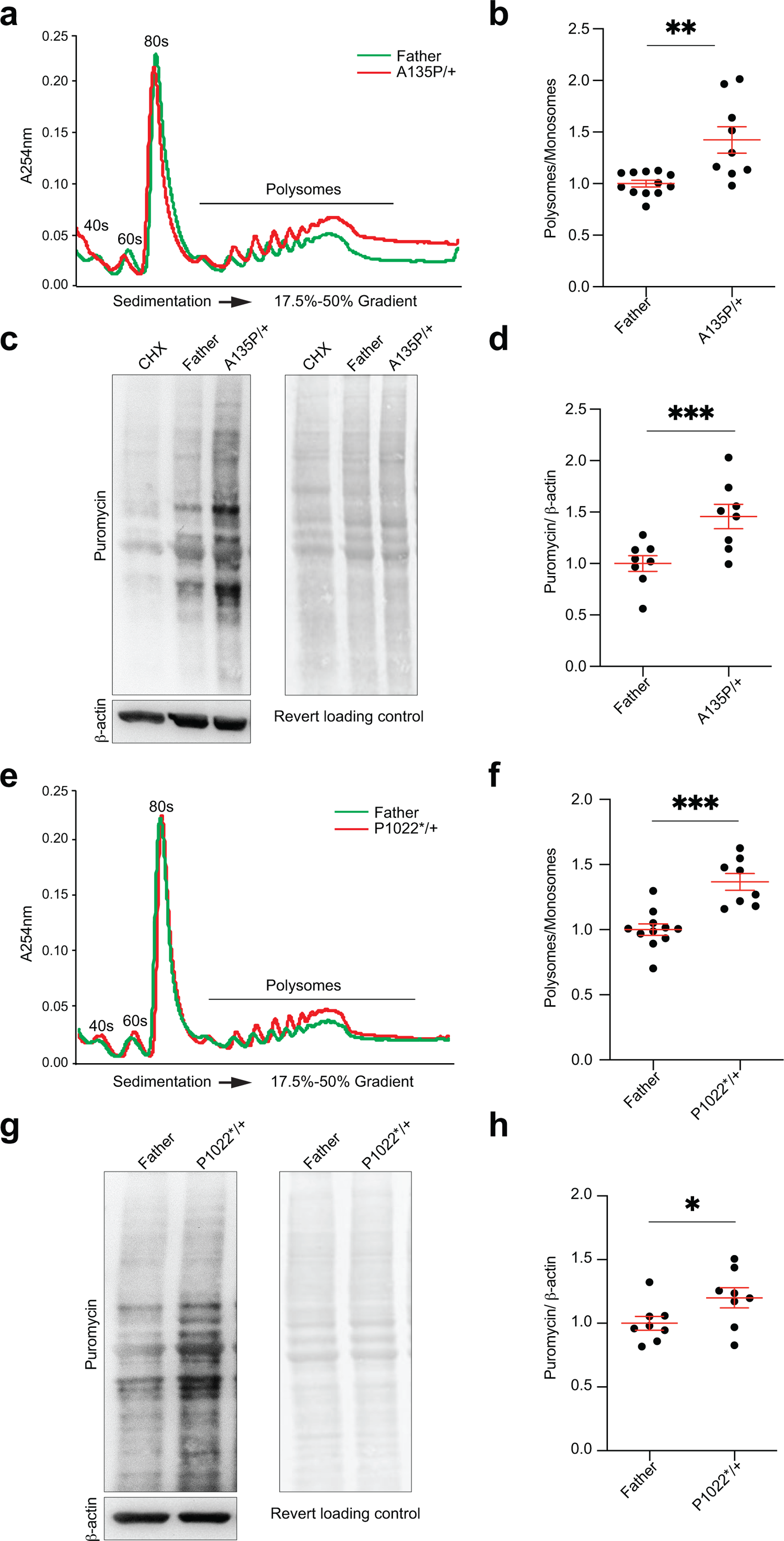
TAOK2 with heterozygous A135P kinase mutation or β-specific P1002* mutation enhances general translation and increases protein synthesis in patient derived LCLs. **(a)** Overlay of LCLs polysome profiles from unaffected father and patient with TAOK2 mutated at A135P in kinase domain shows increase in P/M ratio of LCLs patient profiles. **(b)** Quantifications of the normalized P/M ratios for polysome profiling reveal a statistically significant increase in P/M ratios of LCLs patient profiles. Number of biological replicates of LCLs from patients with TAOK2 mutated at A135P in the kinase domain n=9, and n=12 from unaffected father; **P < 0.01, SEM. error bars, unpaired t-test. **(c)** Immunoblot of protein lysates from patient-derived LCLs with TAOK2 mutated at A135P kinase domain and LCLs from unaffected father analyzed by SUnSET assay shows increased amounts of the newly synthesized puromycin-labelled protein in patient cells. Cycloheximide (CHX) was added to the control cells (father) before being treated with puromycin to verify the specificity of the antibody signal. Revert loading control protein stain was used to show equal protein loading. **(d)** Quantifications of the immunoblot SUnSET assay show a significant increase of densiometric anti-puromycin antibody signal normalized to β-actin in patient-derived LCLs with A135P mutations in the Taok2 kinase domain. Number of puromycin-treated LCLs replicates from unaffected father and patient were 8 per condition; ***P < 0.001, SEM. error bars, unpaired t-test. **(e)** Overlay of LCLs polysome profiles from unaffected father and patient with TAOK2β mutated at P1022* shows an increase in P/M ratio of LCLs patient profiles. **(f)** Quantifications of the normalized P/M ratio for polysome profiling reveal a statistically significant increase in P/M ratios of LCLs patient profiles. Number of biological replicates of LCLs from patients with TAOK2β mutated at P1022* n=8, and n=11 for cells from the unaffected father; ***P < 0.001, SEM. error bars, unpaired t-test. **(g)** Immunoblot of protein lysates from patient-derived LCLs with TAOK2β mutated at P1022* and LCLs from unaffected father analyzed by SUnSET assay shows an increase of the newly synthesized puromycin-labelled protein in patient cells. Revert loading control protein stain was used to show equal protein loading. **(h)** Quantifications of the immunoblot SUnSET assay reveal a significant increase of puromycin densiometric signal normalized to β-actin in patient-derived LCLs with TAOK2β mutated at P1022*. Number of puromycin-treated LCLs replicates from unaffected father and patients with TAOK2β mutated at P1022* were 8 per condition; *P < 0.05, SEM. error bars, unpaired t-test.

**Supplementary Figure 6:**
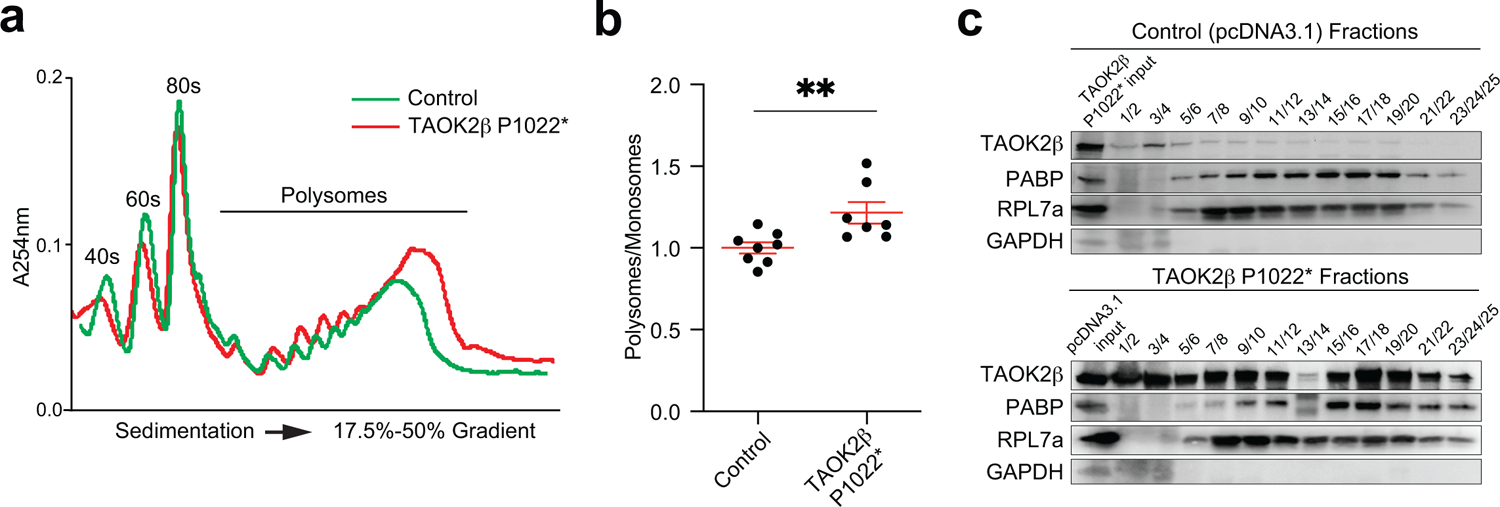
N2a cells expressing TAOK2β P1002* show increased P/M ratios of polysome profiling. **(a)** Overlay of polysome profiles of N2a cells transfected with TAOK2βP1022* or control plasmids shows an increase in the P/M ratio of the profile from cells expressing TAOK2βP1022*. **(b)** Quantifications of the normalized P/M ratios of polysome profiles reveal a statistically significant increase in the P/M ratios for profiles of cells expressing TAOK2βP1022*. Number of biological replicates, TAOK2βP1022* transfected cells n= 7, control n= 8; SEM. error bars, **P < 0.01, unpaired t-test. **(c)** Immunoblots of the pooled gradient polysome fractions of N2a cells either transfected with TAOK2β-P1022* or control plasmids show the presence of TAOK2β-P1022*, detected by TAOK2β antibody, in polysomes of cells expressing TAOK2β-P1022* (lower) compared to the less detected endogenous TAOK2β in polysomes of control non-transfected cells (upper). PABP1 and RPl7a were used as positive controls and GAPDH as a negative control. Input from the total cytoplasmic lysate, either from cells expressing TAOK2β-P1022* or control cells was loaded at the beginning of the blot to verify the transfection efficiency.

**Supplementary Figure 7:**
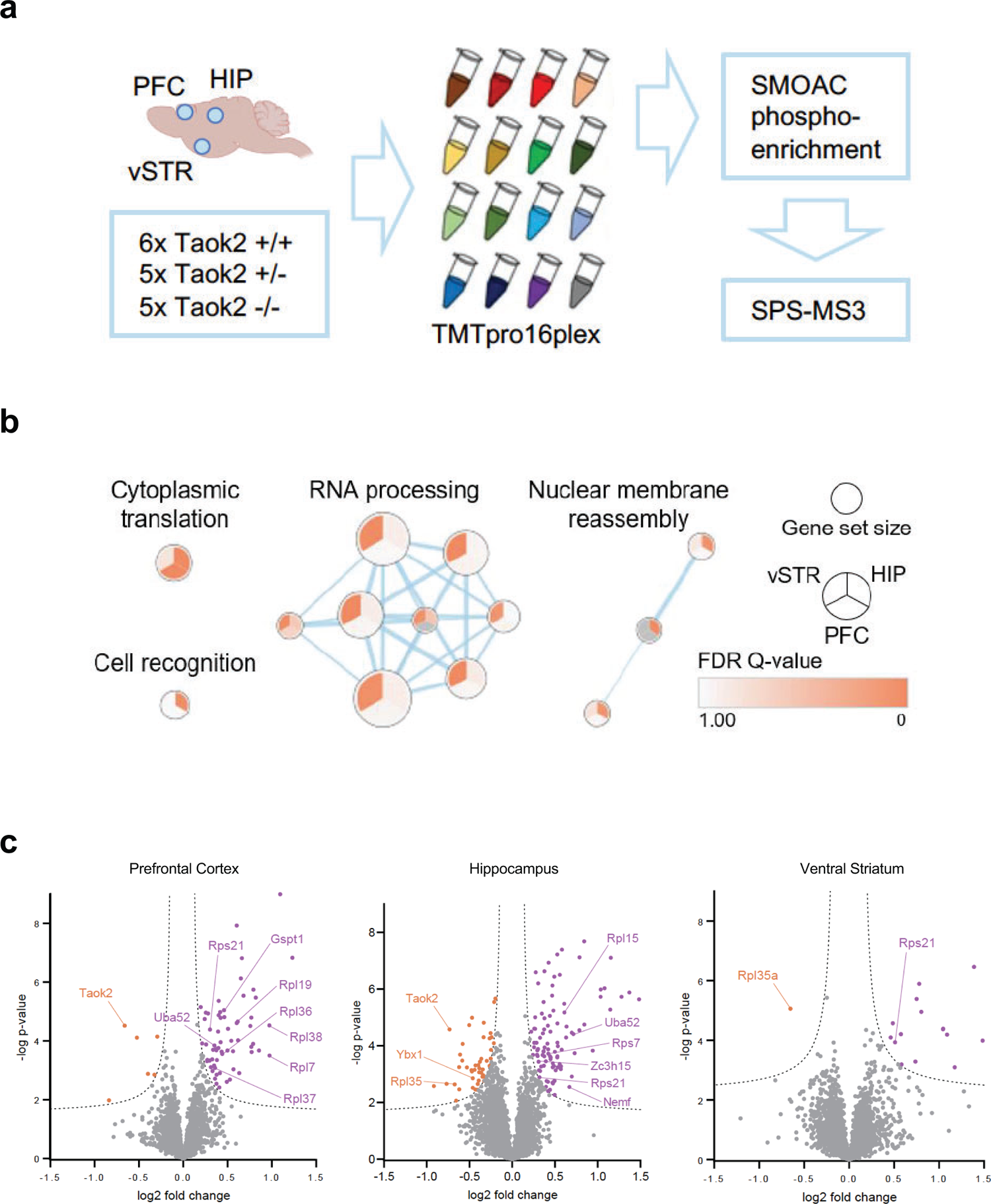
Phosphoproteomics and proteomics of different Taok2 knockout brain regions. **a)** Schematic overview of the phosphoproteomics approach. **b)** Gene set enrichment analysis (GSEA) of pre-ranked Taok2 knockout / Taok2 wild-type total proteomics. The cytoplasmic translation gene set is significantly upregulated in the prefrontal cortex and hippocampus of Taok2 -/- mice. Gene sets with FDR<0.1 for at least one brain region are shown. **c)** Volcano plots of Taok2 knockout / Taok2 wild-type total proteome shows multiple individual proteins involved in cytoplasmic translation are differentially expressed in Taok2-/- brain regions. Purple = significantly upregulated in Taok2 -/-, orange = significantly downregulated in Taok 2-/-. FDR=0.05, S0=0.05. One data point is outside of axis limits.

**Supplementary Figure 8:**
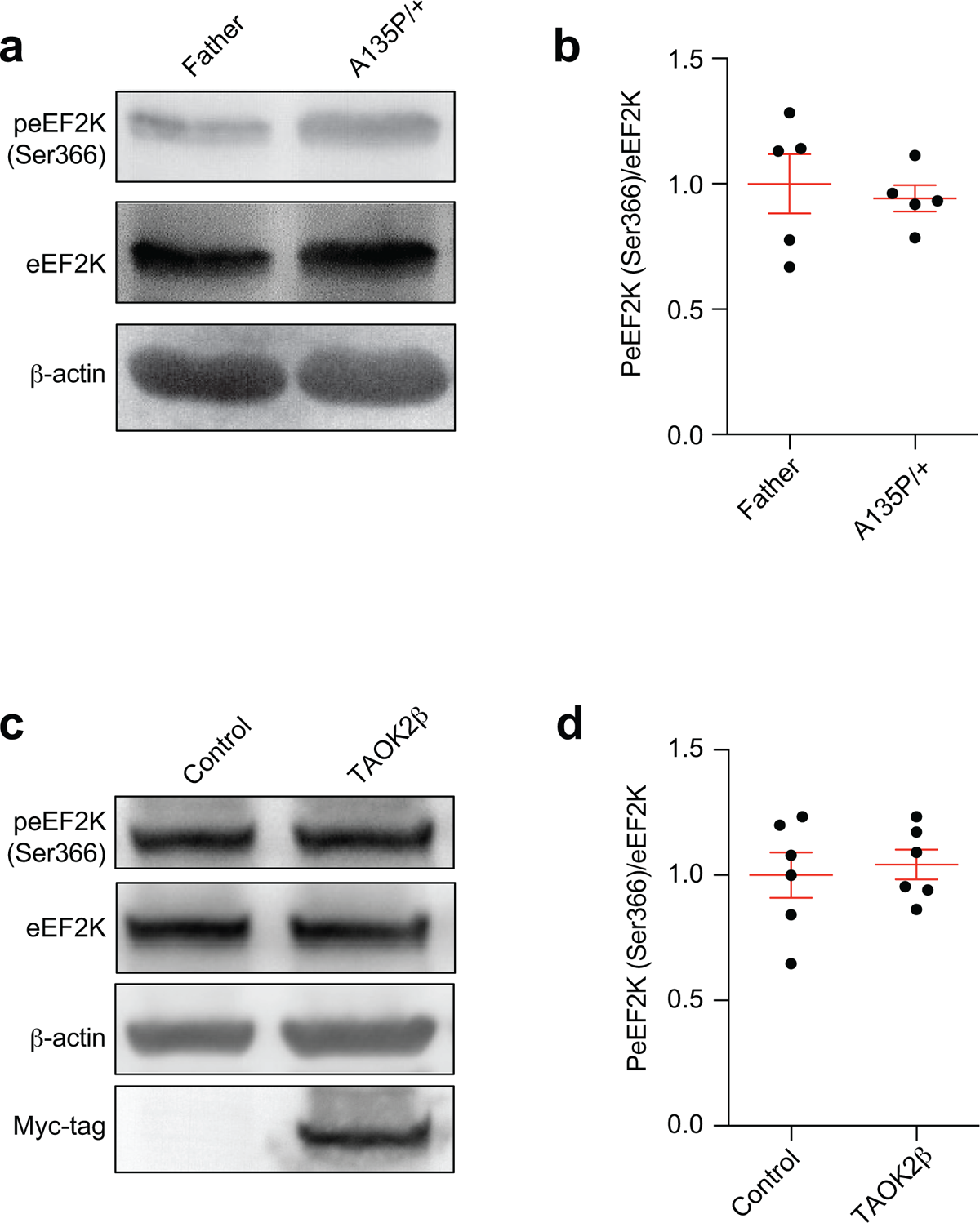
TAOK2 with heterozygous A135P kinase mutation in patient derived LCLs or TAOK2β overexpressed in N2a cells does not affect eEF2K (Ser366) phosphorylation. **(a, b)** Immunoblots and quantifications of protein lysates from LCLs with TAOK2 mutation A135P at its kinase domain and non-affected father show no significant change in the ratio of phospho-eEF2K (Ser366) to total eEF2K. Number of biological replicates were n=5 per condition; unpaired t-test. β-actin was used as a loading control. **(c, d)** Immunoblots and quantifications of protein lysates from N2a cells expressing wild-type TAOK2β and control cells show no significant change in the ratio of phospho-eEF2K (Ser366) to total eEF2K. Number of biological replicates were n=6 per condition from different independent transfection experiments; unpaired t-test. β-actin was used as a loading control. Cells were starved for 24 hours (the usual medium in the study without serum) before harvesting and lysis.

**Supplementary Figure 9:**
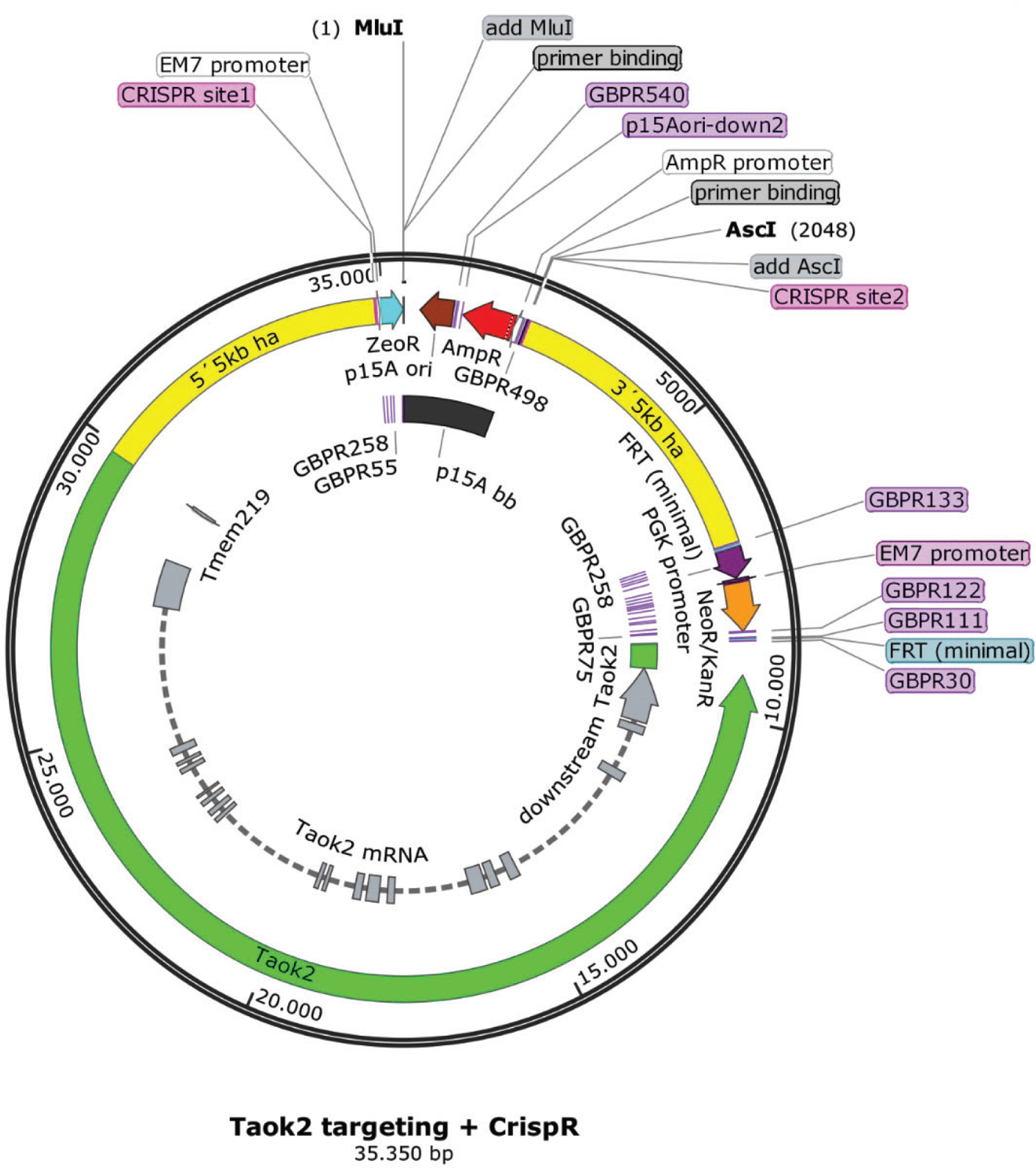
Map for the plasmid RP23-321H3_bb_Taok2 PB3’Amp_PB5’Kan shows the integration of Taok2 by PiggyBac transposase. PiggyBac integration sites flank 21.2 kb murine genomic DNA (green) from the murine chromosome 7: 126,464,301-126,485,468 of GRCm39 reference genome from ENSEMBL covering the murine Taok2 gene.

